# A Low-Dose Triple Antibody Cocktail Against the Ectodomain of the Matrix Protein 2 of Influenza A Is A Universally Effective and Viral Escape Mutant Resistant Therapeutic Agent

**DOI:** 10.1101/2022.04.02.486847

**Authors:** Teha Kim, Lynn Bimler, Sydney L. Ronzulli, Amber Y. Song, Scott K. Johnson, Cheryl A. Jones, S. Mark Tompkins, Silke Paust

**Author notes:** Silke Paust, Ph.D., Department of Immunology and Microbiology, The Scripps Research Institute, Immunology Building 313/114, 10466 North Torrey Pines Road, La Jolla, CA 92037. S. Mark Tompkins, Ph.D., Center for Influenza Disease and Emergence Research, University of Georgia, 119 Carlton Street, Center for Vaccines and Immunology, Rm 1506, Athens, GA 30602. Authorship note: TK and LB are co-first authors.

## Abstract

Influenza, a negative sense single-strand RNA virus of the genus Orthomyxoviridae, causes respiratory illness in humans and animals and significant morbidity and mortality worldwide. While exposure to a specific influenza A strain causes homologous protection, two immune-dominant influenza A virus (IAV)-encoded epitopes - Hemagglutinin (HA) and Neuraminidase (NA) - undergo antigenic shift and drift, resulting in IAVs to which humans lack pre-existing immunity. Without a universal vaccine or therapeutic agent, influenza virus infections will significantly threaten human health. The extracellular domain of the Matrix protein 2-ion channel (M2e) is an ideal antigenic target for a universal influenza therapy: it is highly conserved across influenza A serotypes, has a low mutation rate, and is essential for viral entry and replication. However, less than 20% of humans generate M2e-specific antibodies in response to IAV exposure, thus lacking the benefits of M2e-MAb-mediated immunity. To therapeutically address this deficit, we generated several non-neutralizing M2e-specific monoclonal antibodies (M2e-MAbs) with strong universal IAV treatment potential. Using three MAbs that bind to M2e differentially and competitively, we developed a low-dose M2e-MAb triple cocktail as an effective universal prophylactic and therapeutic agent. We identified the low-dose M2e-MAb triple cocktail’s optimal antibody-clone combination, isotype, minimum effective dosage, and administration time points in mouse models challenged with human and zoonotic BSL-2 and BSL-3 IAV strains, demonstrating its universal potential. Using the IgG2a isotype, which had proven most effective, we established FcγRI, FcγRIII, and FcγRIV as required for M2e-MAb-mediated protection of IAV-challenged mice. Importantly, we established individual M2e-MAbs and the resulting triple cocktail as effective and viral escape mutant-resistant treatments in immunocompetent and immunodeficient mice. These unique qualities provide precedence for prioritizing our M2e-MAbs for therapeutic development.

**CONFLICT OF INTEREST STATEMENT:** SP serves on the scientific advisory board for Shoreline Biosciences, Qihan Biotechnology and is a Scientific Consultant for Qihan Biotechnology and the Genomics Institute of the Novartis Research Foundation. The remaining authors declare no competing interests.

## INTRODUCTION

Influenza A, B, C, and D viruses are lipid-enveloped viruses, and antibody-based immune responses to two of its main membrane proteins, HA and NA, are used to determine influenza A serotypes ^1, 2^. IAVs infect multiple species, including humans, birds, pigs, and other animals, while influenza B and C typically only infect humans ^2^. Exposure to a specific influenza A strain causes robust immunity and homologous protection. However, seasonal variations in influenza A, mostly due to point mutations in HA and NA called “antigenic drift”, result in new IAV strains to which humans lack prior exposure and are thus highly susceptible ^3–5^. Less frequently, co-infections allow genetic HA and NA gene re-assortments in a process called “antigenic shift”, resulting in novel and potentially highly virulent IAV subtypes with pandemic potential ^2^. In addition, localized outbreaks of avian “Highly Pathogenic Avian Influenza A” (HPAI) viruses (H5N1 A/Vietnam/1203/2004 (H5N1), and H7N9 A/Anhui/1/2013 (H7N9)) continue to threaten global public health, with HPAI strains occasionally spilling over into the human population, resulting in a 40-65% mortality rate ^6^.

Influenza virus infections cause about one billion illnesses and half a million deaths worldwide annually ^7^. These deaths include approximately 50,000 persons in the US, where the annual cost to the US economy is an estimated 11.2 billion US dollars ^8^. There are currently six FDA-approved treatments for IAV infection ^9,10^, including NA and M2e inhibitors. However, viral escape mutants have developed to each in clinical trials and/or during seasonal or pandemic outbreaks ^11–19^, and some mutations increase the transmissibility of resistant viruses ^16^. Two of these treatments, amantadine and rimantadine, are M2 channel blockers now rendered ineffective due to widespread resistance caused by M2 mutations preventing their binding but preserving the M2 channel activity^20^. By 2009, all H3N2 and H1N1 isolates tested were resistant to adamantane treatment ^11^. Without an FDA-approved universal influenza virus vaccine, we depend on annual vaccine production, but current vaccination rates for US children and adults are well under 50 percent ^21^. Annual vaccine production depends on strain predictions based on circulating strains; thus, efficacy varies yearly. Once strains have been selected for seasonal vaccine production, it takes at least six months to manufacture and distribute a seasonal influenza virus vaccine ^22^. Consequently, should a pandemic strain develop after seasonal vaccine efforts have begun, vaccine availability can lag peak infection rates by several months, as it did during the most recent H1N1 pandemic in 2009 ^23–26^. Thus, viral resistance to common IAV treatments will be especially detrimental during the time required for pandemic influenza vaccine development.

Effective, universal, escape mutant-resistant “off the shelf” IAV therapeutic agents, such as those we report here, enhance pandemic preparedness. They are critical to protecting human health when seasonal vaccine efficacy is low and offer treatment options to the immunocompromised who do not benefit from vaccination, as well as the unvaccinated. Although HA and NA are highly immunogenic, they are not generally appropriate target antigens for a universal IAV therapeutic or prophylactic agent due to their high mutation rates ^2^. Also, while some broadly protective NA and HA stalk (HA-2) antibodies have been identified, none are established as escape mutant resistant, and none are FDA-approved ^27–29^. The IAV-encoded highly conserved proton channel M2, which is required to disassemble influenza’s viral core, and is essential for viral entry, replication, assembly, and budding ^30–32^, is expressed on influenza virions and infected cells. It’s 5-prime coding region is shared between the M1 and M2 proteins ^25^, the N-terminus is highly conserved across different IAV serotypes ^33–35^. Antibodies specific to this conserved region are more resistant to escape mutants ^36^; however, less than 20% of IAV-infected humans mount a natural M2e-specific antibody response ^37^, perhaps due to the extracellular domains small size and rarity. M2e-MAbs are considered an excellent strategy for a broadly effective IAV therapeutic agent ^32^.

Non-neutralizing antibodies are an essential component of the human immune response to infection. They protect with their Fc-effector functions, such as complement-mediated cytotoxicity (CMC), neutrophil and monocyte phagocytosis activation, and NK cell-mediated antibody-dependent cytotoxicity (ADCC) ^38^. FcγRs are broadly expressed on multiple immune cell types residing in various tissues in mice ^39^. Murine IgG1 binds to the low-affinity inhibitory receptor FcγRIIb (CD32), and the low-affinity activating receptor FcγRIII (CD16), whereas IgG2a binds to the activating receptors FcγRI (CD64), FcγRIII (CD16), and FcγRIV (CD16-2) ^40–42^. The high-affinity activating receptor FcγRI is predominantly expressed by tissue-resident macrophages and myeloid dendritic cells ^43–45^, and the anti-viral cytokine IFNγ can induce its expression. FcγRIII is expressed by myeloid cells (monocytes, macrophages, and dendritic cells), granulocytes (neutrophils, eosinophils, basophils, and mast cells), and NK cells, triggering NK cell-mediated ADCC ^46^, cytokine production, and inflammation ^47, 48^. In contrast, B and T cells do not express FcγRIII ^49^. The more recently discovered FcγRIV is thought to activate monocytes, macrophages, dendritic cells, and neutrophils but is absent on lymphocytes ^42^. The mouse IgG2a isotype, thought to be equivalent to human IgG1 ^39^, preferentially binds to the activating FcγRs, while IgG1 can trigger both activating (FcγRIII) and inhibitory (FcγRIIb) FcγRs ^41^. Therefore, compared to IgG1, murine IgG2a, when administered systemically, is more effective in fighting viral infections, including Ebola ^50^, yellow fever ^51^, and influenza ^52^. However, antibody-based therapies have not yet been approved to treat influenza virus infection in humans. Our impactful study addresses this therapy gap and contributes significantly to the field of IAV-specific immunity and therapeutics development. Also, this study will provide crucial mechanistic insight into how non-neutralizing antibody responses contribute to protection from severe disease.

## RESULTS

### M2e-specific antibodies bind to M2e’s N-terminal region

We recently reported the generation of several murine M2e-MAbs, demonstrated their M2e-antigen specificity and breadth, and established their efficacy as IAV prophylactic agents ^53^. All of our M2e-MAbs recognized M2e peptides encoded by at least eight distinct IAV serotypes and their corresponding influenza A virions and IAV-infected MDCK-ATL cells ^53^. Clones 472 (IgG2a), 522 (IgG1), and 602 (IgG2a) also demonstrated universal potential by significantly improving survival rates and weight loss in highly susceptible Balb/c mice challenged with a lethal dose of PR8, CA07, H5N1, or H7N9 ^53^. In this study, we used an M2e-consensus sequence (CS) alanine scanning peptide library (**Table S1** and **Fig. 1 A**), and an 18-mer overlapping M2e-truncation peptide library (**Fig. 1 B**) to test the binding of biotinylated IgG1 isotype M2e-MAb clones 472, 522, and 602 by ELISA. M2e-MAb binding was abrogated or reduced if alanine mutations were introduced into M2e’s highly conserved N-terminus (**Fig. 1 A**). Truncation mutants lacking the amino acid serine at position 1 also abrogated the binding of all three M2e-MAbs (**Fig. 1 B**). Further, comparing M2e-MAb binding to M2e-peptides with either the original amino acid serine (polar uncharged side chains), lysine (positively charged side chains), aspartic acid (negatively charged side chains), or alanine (hydrophobic uncharged side chains) established that clones 602 and 522 bind to M2e-peptides with a serine in the first position, and clone 472 binds to M2e-peptides with a serine or an alanine in the first position, but none of the clones can bind to M2e-peptides with a charged amino acid in the first position (**Fig. 1 C**). Our data clearly demonstrate that M2e-MAb clones 472, 522, and 602 bind M2e’s highly conserved N-terminal region, requiring a serine (clones 472, 522, or 602) or an alanine (clone 472) at position 1. As the N-terminus of the M2e-protein is highly conserved between IAV serotypes, our data explain why M2e-MAb clones 472, 522, and 602 bind broadly to various IAV serotypes ^53, 54^, and substantiate their strong potential as a universal IAV treatment.

**Figure 1.**
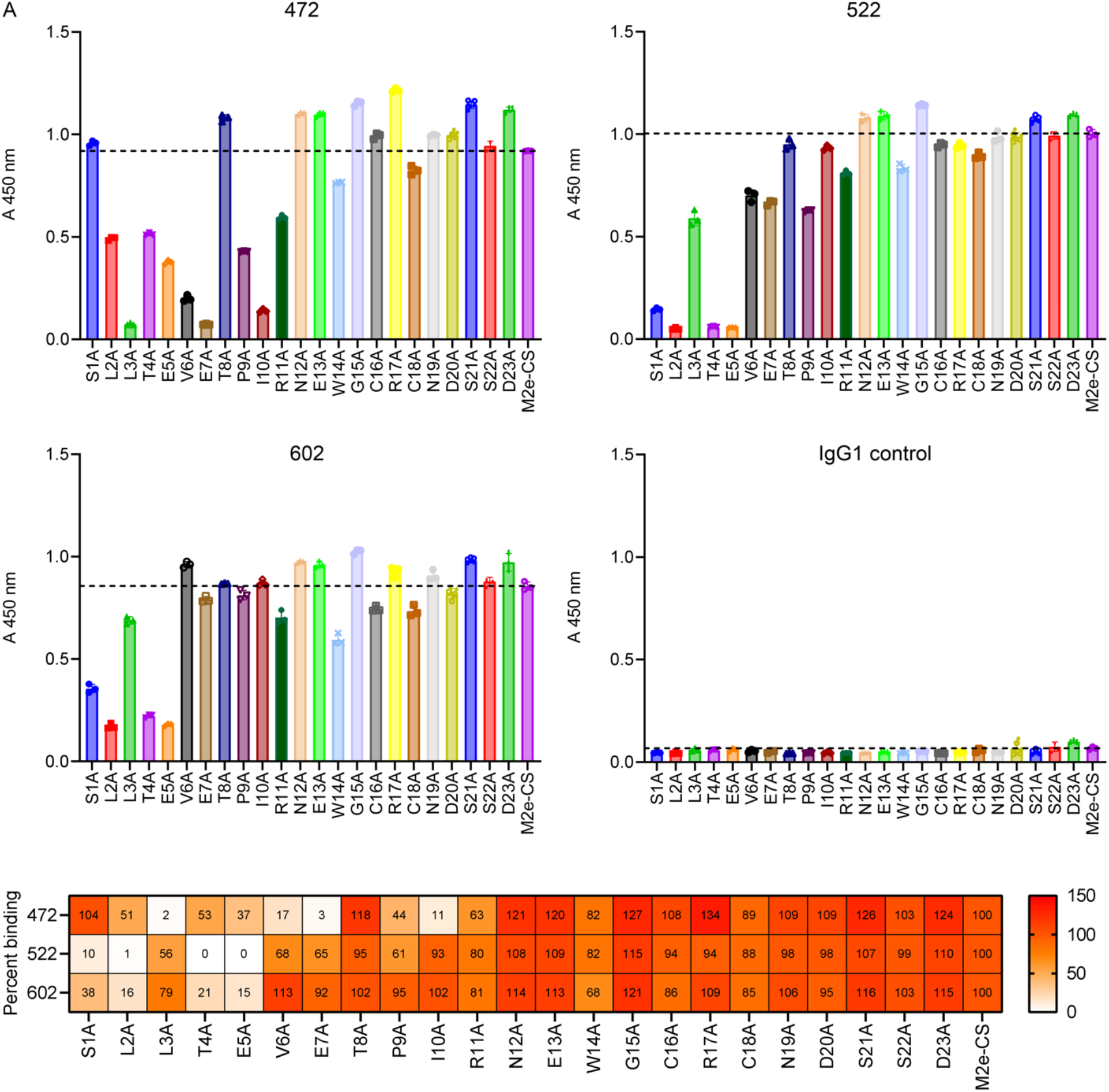
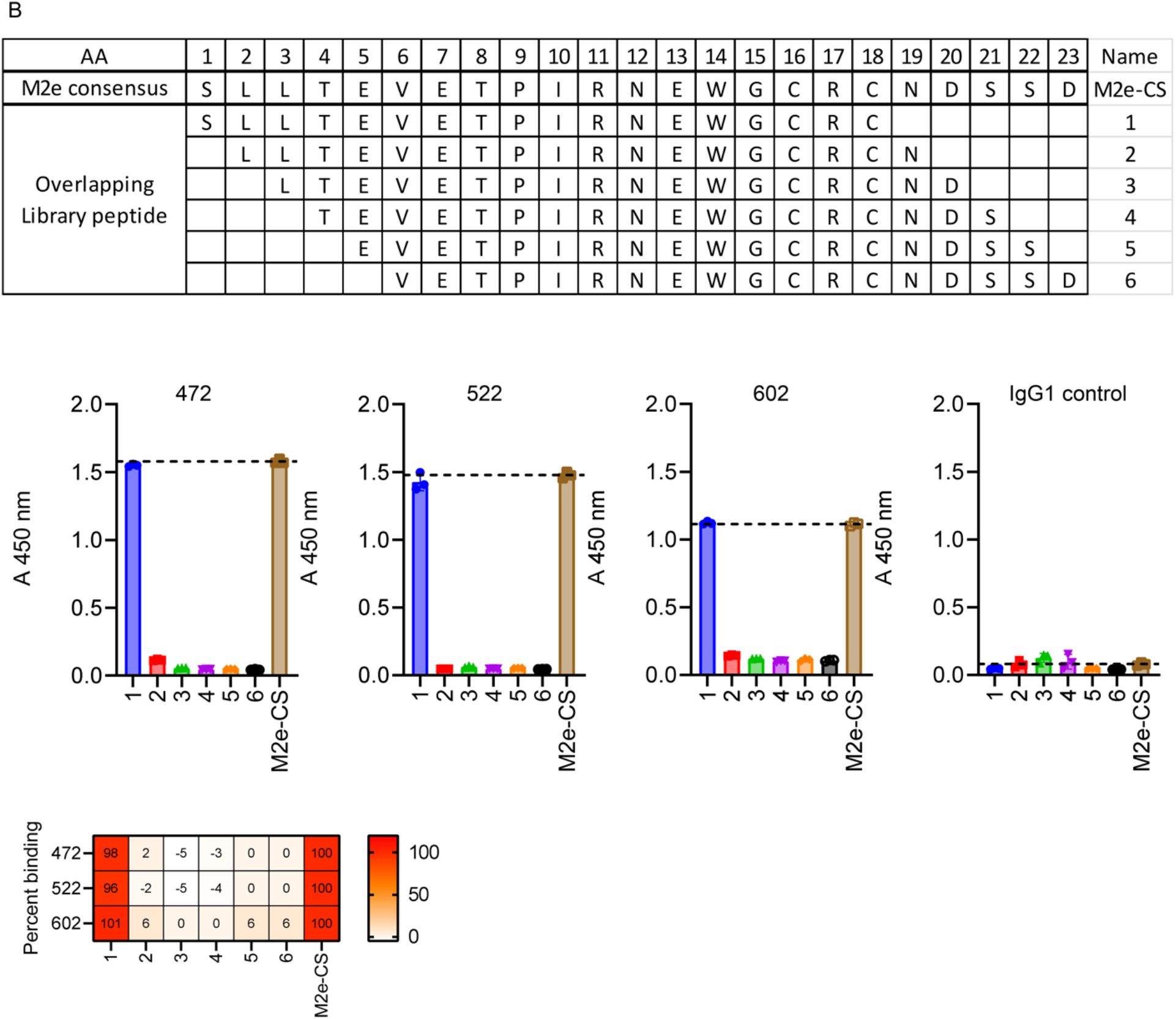
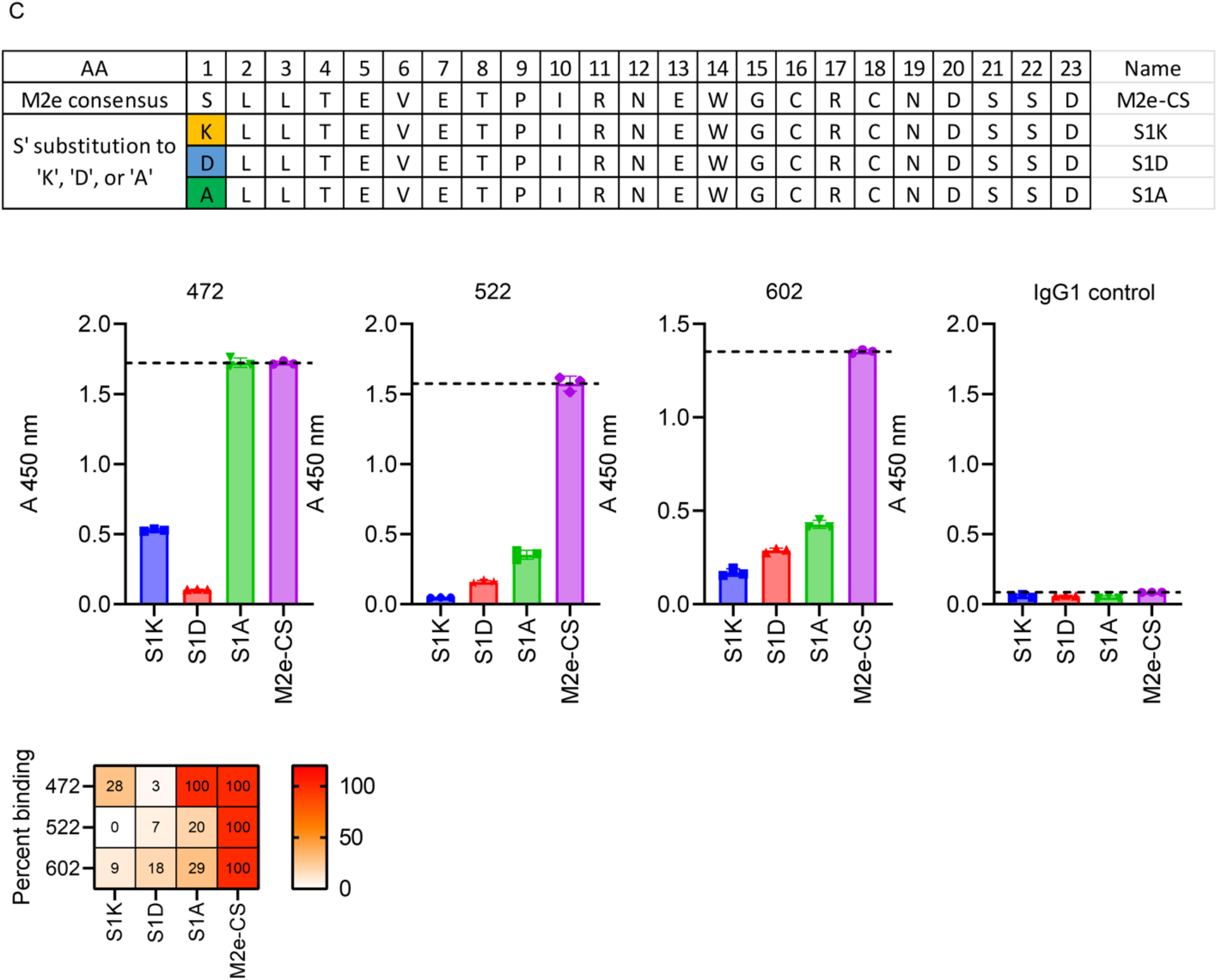
Clones 472, 522, and 602 bind M2e in M2e’s highly conserved N-terminal region. Clones 472, 522, 602, all expressed as IgG1 isotypes, and IgG1 isotype-matched control MAb were biotinylated for epitope mapping by ELISA. Corning^®^ 96-well EIA/RIA assay plates were coated with 2.5 µg/ml of either (**A**) the M2e-consensus sequence (CS) peptide and corresponding alanine scanning peptides, (**B**) 18-mers of M2e-CS overlapping peptides, or (**C**) opposite or neutral charge M2e-CS peptides, in which the first amino acid Serine (S)* was substituted with Lysine (K)^†^, Aspartic Acid (D)^‡^, or Alanine (A)^§^, as indicated, to confirm a requirement for the first amino acid (serine) for M2e-MAb binding. Serine (S) has a polar uncharged side chain, Lysine (K) has a positively charged side chain, Aspartic Acid (D) has a negatively charged side chain, and Alanine (A) has a hydrophobic side chain. Then, 2.5 µg/ml of the specified M2e-MAb clone was used to determine the clone’s binding to the indicated peptide by ELISA. To calculate the percent binding for the heatmaps, the average binding - as determined by the absorbance at 450 nm (A 450nm) of each M2e-MAb clone and with the isotype-matched control antibody-determined background subtracted - was used to calculate the percent binding as related to the consensus peptide sequence control, which was set as 100 percent.

### M2e-specific antibodies bind to M2e competitively

Our data reveal that the M2e-binding sites for clones 472, 522, and 602 are similar but not identical (**Fig. 1 A**). Thus, we used inactivated influenza virions to perform competition assays to determine if M2e-MAb clones 422, 572, and 622 compete for binding. Competitive binding to the following IAV serotypes was evaluated: H1N1 A/PR/8/34 (PR8), pH1N1 A/CA/07/2009 (CA07), A/Vietnam/1203/2004 (VN1203), A/Anhui/1/2013 (Anhui1) (**Table 1 and Fig. 2**), and A/FM/1/1947 (FM1), A/sw/NE/A01444614/2013 (swNE), A/sw/TX/A01049914/2011 (swTX), and A/sw/MO/ A01444664/2013 (swMO) (**Table 1** and **Fig. S1**). We added M2e-MAbs sequentially to plates coated with the indicated inactivated virions; first, an unlabeled competitor M2e-MAb and then a biotinylated monitored antibody for detection by ELISA. In this assay, a reduction of the detected absorbance indicates interference by the unlabeled competing antibody with the monitored antibody’s binding. As expected, generally, a given antibody clone strongly competes with itself **(Fig. 2)**. However, despite the clones similar binding capacity across the different IAV serotypes, some competition was observed: The recognition of PR8 and CA07 (**Fig. 2, A, B, and E**) by clones 472, 522, and 602 was more similar than that of VN1203 and Anhui1 (**Fig. 2, C, D, and E**), where clone 522 outcompeted clone 602 with a higher affinity for a shared VN1203-M2e epitope (**Fig. 2, C and E**). Generally, clone 522 was most able to compete with the other clones (**Fig. 2 E** and **Fig. S1 E**). Similar results were obtained when we tested various contemporary IAV strains (**Fig. S1**). Our data demonstrate that M2e-MAb clones 472, 522, and 602 recognize overlapping epitopes in the N-terminus of the highly conserved M2e-protein resulting in their differential but competitive binding. These findings are consistent with our previously published results ^53^ establishing broad but distinct M2e-MAb clones 472, 522, and 602 mediated protection against many IAV serotypes with serotype-specific M2e-sequences.

**Figure 2.**
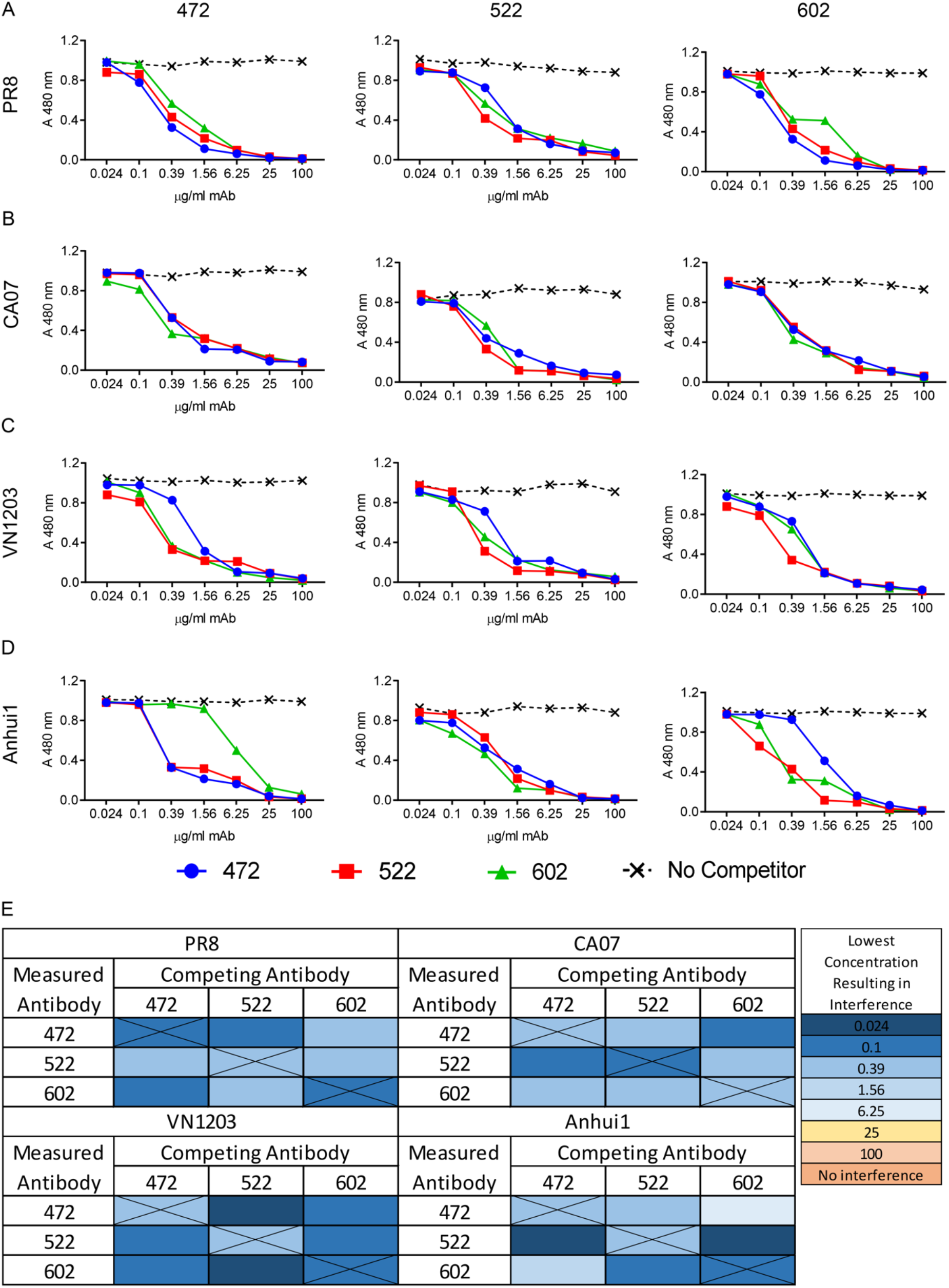
M2e-specific antibodies bind to M2e competitively. Inactivated virions from (**A**) PR8, CA07, (**C**) VN1203, and (**D**) Anhui1 were used as coating antigens to determine the competitive binding of biotinylated M2e-MAb clones 472 (IgG2a), 522 (IgG1), and 602 (IgG2a) (2 μg/ml) by competition ELISA. The competing antibody was added before the biotinylated antibody at 4-fold dilutions starting at 100 μg/ml. Absorbance was measured with a biotin-binding secondary antibody. (**E**) The concentration at which the absorbance dropped 0.1 below the average absorbance of the “no competitor” control (specific to the antibody and virus) was used to summarize the data.

**Table 1.** M2e sequences of the influenza A viruses used in this study and compared to the M2e consensus sequence. Indicated M2e sequences of Influenza A viruses, isolated from humans, birds, and swine ^91^. The consensus sequence is derived from seasonal influenzas viruses circulating since 1957 (H1N1, H2N2, and H3N2; shown). Blue highlights identify mutations in influenza A virus serotypes compared to the consensus sequence.

| Virus | Subtype | Strain Abbreviations | Amino Acid Sequence |  |  |  |  |  |  |  |  |  |  |  |  |  |  |  |  |  |  |  |  |  |  |
| --- | --- | --- | --- | --- | --- | --- | --- | --- | --- | --- | --- | --- | --- | --- | --- | --- | --- | --- | --- | --- | --- | --- | --- | --- | --- |
|  |  |  | 1 | 2 | 3 | 4 | 5 | 6 | 7 | 8 | 9 | 10 | 11 | 12 | 13 | 14 | 15 | 16 | 17 | 18 | 19 | 20 | 21 | 22 | 23 |
| Consensus sequence |  |  | S | L | L | T | E | V | E | T | P | I | R | N | E | W | G | C | R | C | N | D | S | S | D |
| A/PR/8/1934 | H1N1 | PR8 | S | L | L | T | E | V | E | T | P | I | R | N | E | W | G | C | R | C | N | G | S | S | D |
| A/CA/07/2009 | pH1N1 | CA07 | S | L | L | T | E | V | E | T | P | T | R | S | E | W | E | C | R | C | S | D | S | S | D |
| A/FM/1/1947 (WT & MA) | H1N1 | FM1 | S | L | L | T | E | V | E | T | P | T | K | N | E | W | E | C | R | C | S | D | S | S | D |
| A/Vietnam/1203 /2004 | H5N1 | VN1203 | S | L | L | T | E | V | E | T | P | T | R | N | E | W | E | C | R | C | S | D | S | S | D |
| A/Anhui/1/2013 | H7N9 | Anhui1 | S | L | L | T | E | V | E | T | P | T | R | T | G | W | E | C | N | C | S | G | S | S | E |
| A/sw/NE/A01444614/2013 | H1N1 | swNE | S | L | L | T | E | V | E | T | P | T | R | N | G | W | E | C | K | C | N | D | S | S | D |
| A/sw/TX/A01049914/2011 | H3N2 | swTX | S | L | L | T | E | V | E | T | P | T | R | S | E | W | E | C | R | C | S | D | S | S | D |
| A/sw/MO/A01444664/2013 | H1N2 | swMO | S | L | L | T | E | V | E | T | P | T | R | N | G | W | E | C | K | C | N | D | S | S | D |

### M2e-MAbs are more protective against lethal IAV challenge as a triple cocktail

We previously established that prophylactic therapy with a cocktail comprised of the two or three most effective M2e-MAbs (clones 472, 522, and 602), protects Balb/c mice from lethality upon PR8, CA07, VN1203, and Anhui1 challenge ^54^. We next experimentally test our hypothesis that an M2e-MAb cocktail ensures therapeutic efficacy at a lower dose than single antibody treatments. To do so, we prophylactically treated Balb/c mice with a 30 μg dose of either the M2e-MAb triple cocktail (10 μg each of clones 472 (IgG2a), 522 (IgG1), and 602 (IgG2a)), or an M2e-MAb double cocktail comprised of either 15 μg each of clones 472 and 522, clones 472 and 602, or clones 522 and 602. Upon lethal PR8 challenge, the protection afforded by 30 µg of the M2e-MAb triple cocktail was significantly better (88%) than the protection observed with any combination of two M2e-MAbs (33-44%, **Fig. 3 A**) or individual M2e-MAb cocktail components (published in ^53^). A 60 µg dose of the triple M2e-MAb cocktail also sufficed to significantly protect of Balb/c mice challenged with either laboratory (**Fig. 3 B**) or pandemic IAV strains (**Fig. 3, C-E**). In addition, the M2e-MAb triple cocktail significantly protected Balb/c mice from weight loss upon lethal IAV challenge (**Fig. 3, A-E**). Our results show that the protection provided by the M2e-MAb triple cocktail is the sum of its individual cross-protective M2e-MAb clones rather than attributable to a single MAb or the protective effects of two together. We conclude that, at low doses, cross-protective M2e-MAbs are more protective against lethal IAV challenge as a triple M2e-MAb cocktail. Consistent with our previous report ^53^, where individual M2e-MAb clones 472, 522, and 602 generally did not demonstrate neutralizing activity *in vivo*, the triple M2e-MAb cocktail only modestly decreased lung viral titers in IAV-infected mice, with significant decreases observed only in PR8 and VM1203-challenged animals (**Fig. 3 F**).

**Figure 3.**
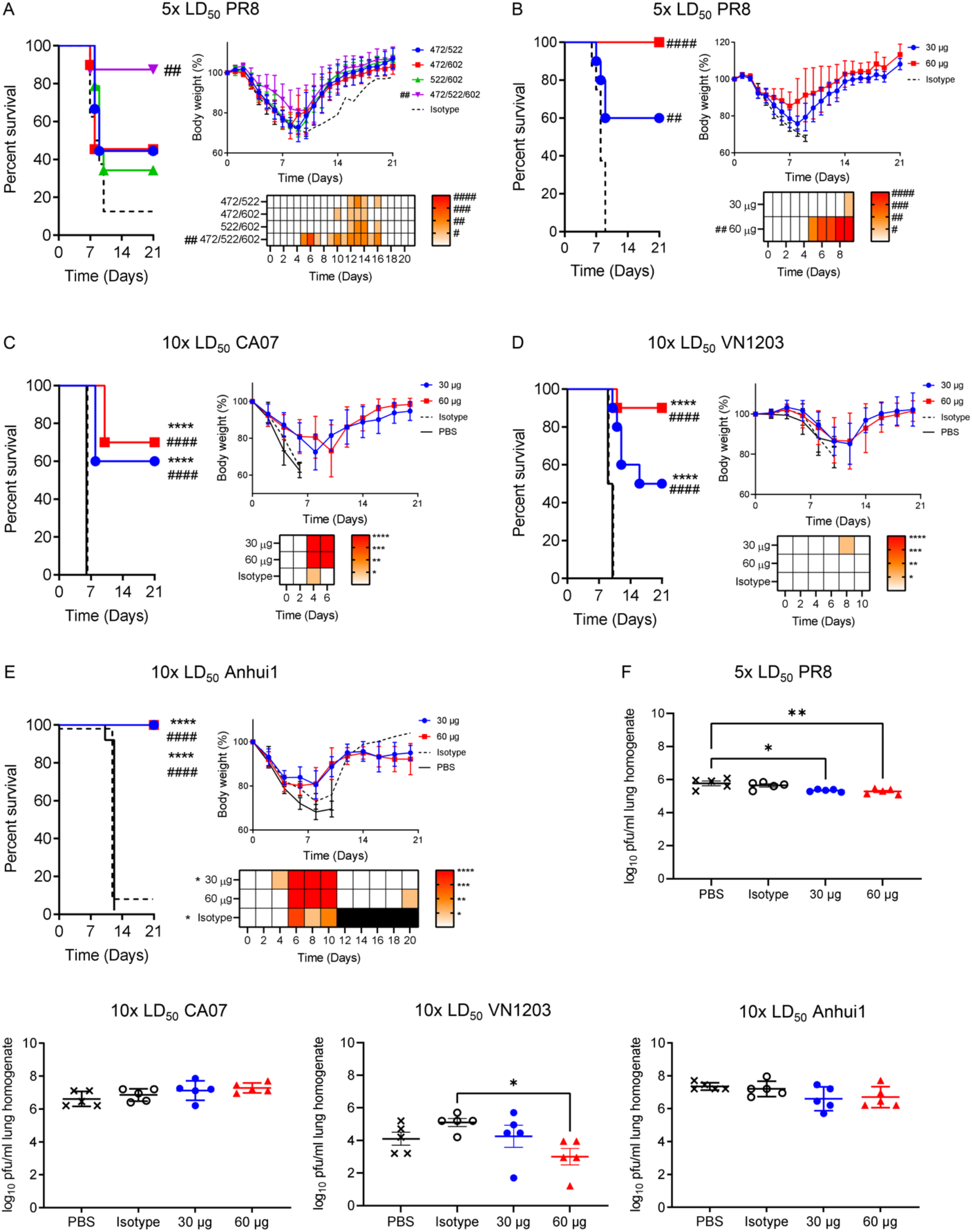
M2e-MAbs are more protective against lethal influenza virus challenge as a cocktail. (**A**) Balb/c mice were treated with a 30 μg dose of the indicated M2e-MAb cocktail (clones 472 (IgG2a), 522 (IgG1), and 602 (IgG2a)), containing two or three M2e-specific antibodies in equal parts, one day before infection with a lethal dose (5x LD50) of PR8. (**B-E**) Balb/c mice were treated with the indicated dose of the M2e-MAb clones 472/522/602 triple cocktail (clones 472 (IgG2a), 522 (IgG1), and 602 (IgG2a)) one day before infection with (**B**) PR8, CA07, (**D**) VN1203, or (**E**) Anhui1. (**A-E**) Percent survival and percent weight loss were recorded. Significant differences in the percent weight loss of the experimental groups compared to their isotype control groups are shown in the heatmap. (**A**) N=8-9, (**B-E**) N=8-10 mice per group. Log-rank (Mantel-Cox) test for survival, and one-or two-way ANOVA (Dunnett’s multiple comparisons) test for percent weight loss. **** or ^####^ p<0.0001, *** or ^###^ p<0.001, ** or ^##^ p<0.01, * or ^#^ p<0.05, with * indicating significance compared to PBS control, and # indicating significance compared to isotype control. Black squares in the heat map indicate the death of the control group animals; thus, no further statistical evaluations could be performed. (**F**) Balb/c mice were treated with the indicated dose of the M2e-MAb triple cocktail one day before infection with PR8, CA07, VN1203, or Anhui1. Lungs were removed on day three post-infection, and viral titers were measured via plaque assay. N=5 mice, **** p<0.0001, *** p<0.001, ** p<0.01, * p<0.05, one-way ANOVA with a Tukey’s multiple comparison test.

### Viral escape mutants do not develop in immunocompetent or immunodeficient mice after viral passaging in the presence of single, cocktail, or alternating M2e-MAb treatments

Viral escape mutants have been reported to all FDA-approved influenza virus therapies, including those targeting M2 function ^55^. Discouragingly, in humans, escape mutants to influenza therapies have arisen as early as 48 hours after treatment ^11, 14^. To determine whether our M2e-MAb clones drive the development of viral escape mutants when administered to IAV-challenged mice, we first passaged PR8 in the presence of the M2e-MAb triple cocktail or PBS (control) in immunocompetent (Balb/c) mice seven times for a total of three and a half weeks (**Fig. 4 A**). Then, we isolated viral RNA from the mice’s lungs to generate influenza M gene segment-specific cDNA, which was subjected to Sanger sequencing and compared to the M2-sequence of the original (day 0) PR8 virus (**Fig. 4 A**). No mutations arose in the M gene in any of our treatment groups despite constant selective and immune pressure from M2e-MAb treatments (**Fig. 4 B**). In addition, we evaluated if single M2e-MAb treatments or alternating M2e-MAb treatments resulted in M-region mutations. Interestingly, we detected no M-region mutations in PR8 preparations isolated after 24 days of single or alternating M2e-MAb treatments (**Fig. S2**). However, mutations outside of the M region can enable viral escape, for example, by delaying M2e-expression ^56, 57^. To exclude this possibility, we challenged Balb/c mice with a lethal dose of the PR8 virus preparation isolated from M2e-MAb triple cocktail treated Balb/c mice, or with control virus isolated from PBS-exposed (control) mice and prophylactically treated one-half of the mice in each group with the M2e-MAb triple cocktail therapy. Notably, the M2e-MAb-triple cocktail maintained robust effectiveness against both the PBS-passaged and the M2e-MAb-triple cocktail-passaged virus (**Fig. 4 C**). These data demonstrate that therapy with individual M2e-MAb clones or the triple cocktail does not result in viral immune escape. Altogether, our sequencing and *in vivo* data demonstrate the virus’s failure to escape from our highly effective antiviral therapies. In contrast to our M2e-MAb therapeutic, viral escape mutants have developed to each of the FDA-approved treatments for IAV infection ^9, 10^ in clinical trials and/or during seasonal or pandemic outbreaks ^11-19^.

**Figure 4.**
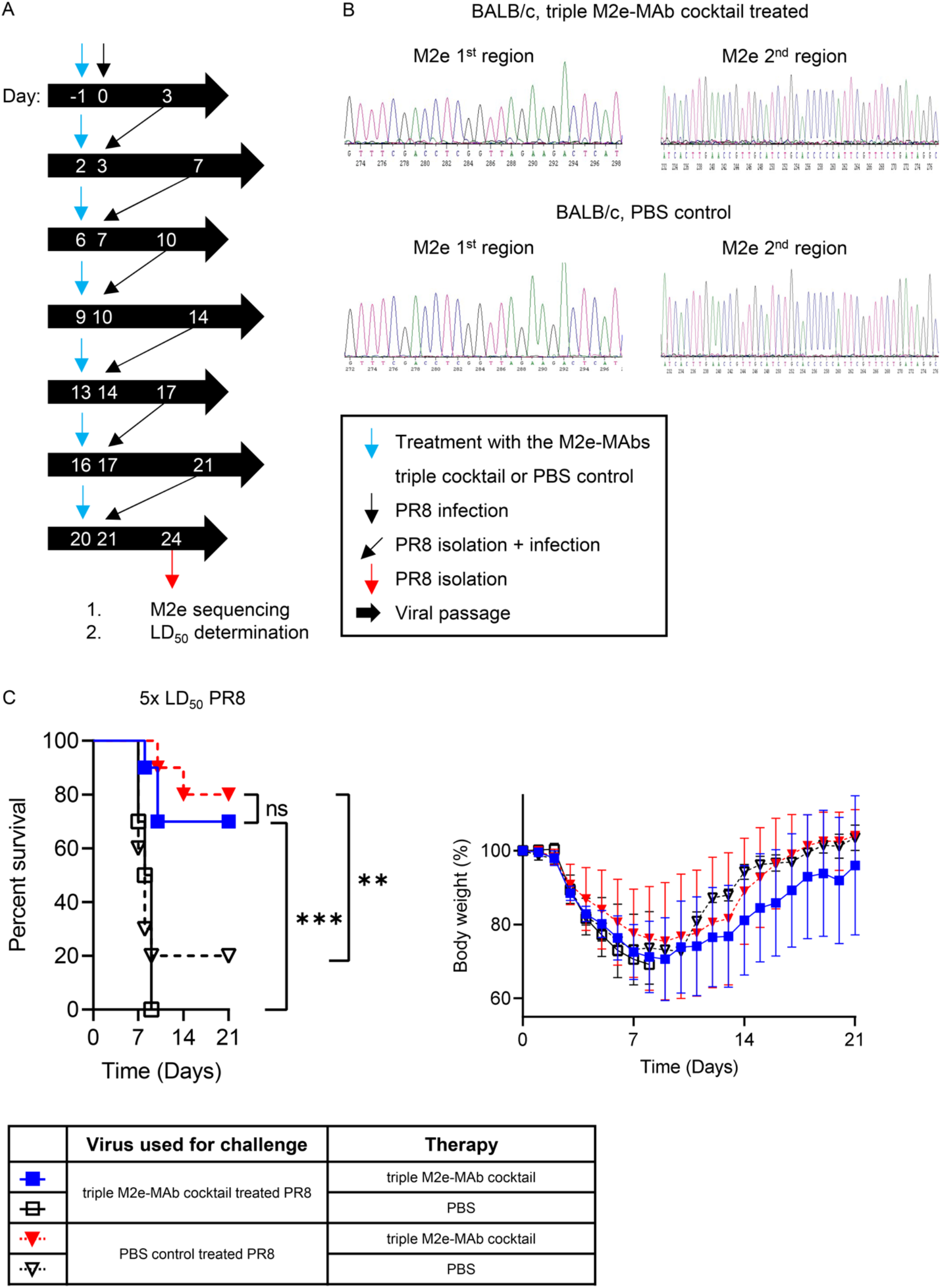
M2e-MAb triple cocktail therapy does not drive the development of viral escape mutants. PR8 “stock virus” was passaged through WT mice for 24 days, and the final viral isolates were analyzed by Sanger sequencing. (**A**) Outline of the time points for mouse-to-mouse passaging of lung-PR8-isolates and the indicated M2e-MAb triple cocktail treatments (clones 472 (IgG2a), 522 (IgG1), and 602 (IgG2a)) in wild-type (WT) mice. At each passage, virus was isolated from lung homogenates of M2e-MAb triple cocktail or PBS control-treated mice and used to infect a group of naïve prophylactically M2e-MAb triple cocktail therapy or PBS (control) treated groups of mice. Intraperitoneal (IP) injections of the M2e-MAb triple cocktail or PBS (control) are indicated by a blue arrow. (**B**) Sequencing chromatograms from the viral isolates isolated from therapeutically or control PBS-treated groups of WT mice. (**C**) Balb/c mice were treated with 60 μg of the M2e-MAb triple cocktail (clones 472 (IgG2a), 522 (IgG1), and 602 (IgG2a)) one day before infection with a lethal dose of (5x LD_50_) of PR8 that had been isolated from either M2e-MAb triple cocktail treated or PBS control treated WT mice. Then, their survival and percent weight loss was determined. N=10 mice/group. Log-rank analysis (Mantel-Cox) test for survival, **** p<0.0001, *** p<0.001, ** p<0.01, * p<0.05.

Some of the FDA-approved treatments for IAV infection increase the transmissibility of resistant viruses ^16^. Thus, we examined whether M2e-MAb therapy modulates IAV virulence. We found that passaging PR8 in wild-type mice in the presence of either individual M2e-MAbs or the triple M2e-MAb cocktail lowers its virulence, as twice as many pfu were needed to reach an LD50 when the virus was derived from the lungs of mice treated with the triple M2e-MAb cocktail (6 pfu) compared to isotype-matched control MAbs or PBS treated IAV infected mice (3 pfu) (**Fig. S2 E**). Single M2e-MAb treatments revealed that clone 472 most robustly reduced viral fitness (12.7 pfu = LD50), followed by clones 602 and 522 (9.4 and 4.3 pfu = LD50, respectively). Thus, during a seasonal or pandemic outbreak, M2e-MAb therapy may reduce virulence resulting in lower transmissibility and reduced viral persistence.

Severely immunocompromised persons comprise about ∼3% of the US population and are at high risk of substantial influenza-related morbidity and mortality ^58^. Similarly, Recombinase Activating Gene 2 knock-out (*Rag2*-KO) mice, which lack T and B cells but have innate immune cells and NK cells capable of FcR-mediated immunity, develop chronic or fatal influenza virus infections^59, 60^ and rapidly develop viral escape mutants to therapies, including to M2e-MAb treatments ^11, 61^.

To determine if the M2e-MAb triple cocktail ameliorates disease in immunocompromised hosts, we prophylactically and therapeutically treated PR8 challenged *Rag2*-KO mice with either individual M2e-MAbs, alternating M2e-MAb treatments, or the triple M2e-MAb cocktail (**Fig. S3**). Individual treatments with M2e-MAb clones 472, 602, but not 522, alternating single M2e-MAb treatments, and M2e-MAb triple cocktail therapy significantly improved the survival and ameliorated infection-induced weight loss of PR8-challenged *Rag2*-KO mice (**Fig. S3**). Also, comparisons of PR8’s M-gene sequences revealed that no mutant escape viruses developed in PR8-infected Rag-2-KO mice, regardless of the therapy regiment (**Fig. S2, C and D**, and **Table S2**). Our data demonstrate that M2e-MAb therapy ameliorates disease in IAV-infected immunocompromised mice and does not elicit viral escape mutants. Our findings are in stark contrast to previously published M2e-MAbs and FDA-approved M2 inhibitors, which rapidly elicit escape mutants in WT and immunocompromised mice ^57, 61–63^.

### M2e-MAb therapeutic efficacy depends on FcγRI (CD64), FcγRIII (CD16), and FcγRIV (CD16-2)

When expressed as IgG1 or IgG2a isotypes, our M2e-MAbs do not fully neutralize IAV *in vivo* (**Fig. 3 F**), suggesting that FcR-mediated immune functions are responsible for the observed therapeutic effects. In mice, IgG1 and IgG2a isotypes robustly activate FcR-mediated immunity:

IgG2a binds to three activating receptors: FcγRI (CD64), FcγRIII (CD16), and FcγRIV (CD16-2) ^40–42^, however, murine IgG1 also binds to the low-affinity inhibitory receptor FcγRIIb (CD32) and the low-affinity activating receptor FcγRIII (CD16). We recently demonstrated that M2e-MAb clones of both the IgG2a (clones 472 and 602) and IgG1 (clone 522) isotypes robustly protect Balb/c mice from circulating and pandemic IAV challenges ^53^. Therefore, we compared the prophylactic efficacy of the M2e-MAb triple cocktail and its individual component antibodies when expressed as either the IgG1 or IgG2a isotypes (**Fig. S4**). We found individual IgG2a-M2e-MAb clones more effective as a prophylactic therapy at any dose than the matching IgG1-M2e-MAb (**Fig. 5, A-D**). Similarly, while IgG2a M2e-MAb triple cocktail therapy robustly ameliorated weight loss and prevented death in PR8-challenged Balb/c mice, the IgG1 M2e-MAb triple cocktail therapy was largely ineffective (**Fig. 5 E**).

**Figure 5.**
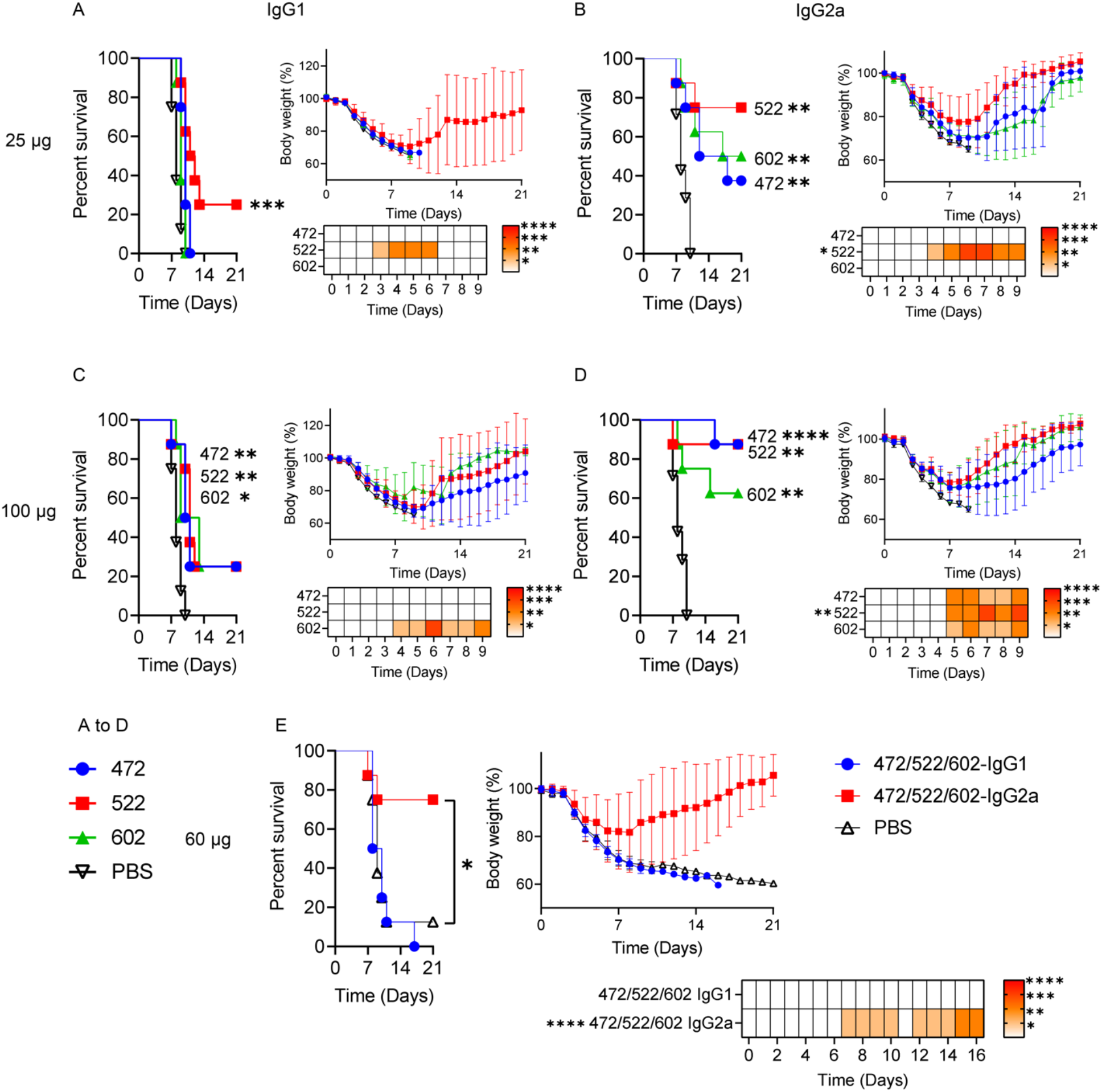
M2e-MAbs expressed as IgG2a isotypes are more protective than IgG1 isotypes. (**A-E**) At -1 dpi, 6–8-week-old female Balb/c mice were infused intraperitoneally with (**A, B**) 25 µg, (**C, D**) 100 µg of the specified M2e-MAbs, or (**E**) 60 µg (each for 20 µg) of the indicated triple cocktail. At 0 dpi, the mice were infected intranasally with a lethal dose (5x LD50) of H1N1 A/PR/8/34. Survival and weight loss were monitored for 21 dpi. N=7-8. * p < 0.05, ** p < 0.01, *** p < 0.001, and **** p < 0.0001, log-rank (Mantel-Cox) test for survival and one- or two-way ANOVA (Dunnett’s multiple comparisons test for percent weight loss. Percent weight data for survival analysis is shown to the right of each graph. The heatmap below each weight loss curve indicates significantly different percent weight from the control group on each day. For heatmap, * indicates significance compared to PBS control. (**A**-**D**) 25 µg and 100 µg data are displayed as two sets for clarity with the shared PBS group.

Based on these data, to identify the relevant IgG2a-dependent FcR mediated effector functions, we engineered the IgG2a-isotype of clone 602 to carry the following Fc mutations: “LALA-PG”, which does not bind to any FcγRs nor C1q ^64^; “LEEA2KA” which binds to FcγRIIb, FcγRIII, and FcγRIV, but not to FcγRI nor C1q ^65^; and “L235E”, which abrogates binding to FcγRI ^65^, and will distinguish between FcγRI and C1q binding (**Fig. 6, A and B**). Our detailed mechanistic studies revealed that 50% of our IgG2a-M2e-MAbs protective function are mediated by FcγRI, but not through complement activation. Surprisingly, in addition to FcγRI, both FcγRIII (clone: 275003) and FcγRIV (clone: 9E9) ^40^ also contributed to maximal therapeutic efficacy (**Fig. 6, C-E**). Of note, we chose to use blocking antibody administrations (MAb clone 275003 to block FcγRIII and MAb clone 9E9 to block FcγRIV ^40^) to determine their potential roles in M2e-MAb-triple cocktail mediated protection, as no effector function blocking mutations have been identified for FcγRIII or FcγRIV, and as a genetic deficiency in either FcR results in the significant upregulation of the other FcR in mice, making results difficult to interpret ^66^. Altogether, our data demonstrate that protection from IAV lethality by our M2e-MAb clones requires the joint effector functions of FcγRI, FcγRIII, and FcγRIV.

**Figure 6.**
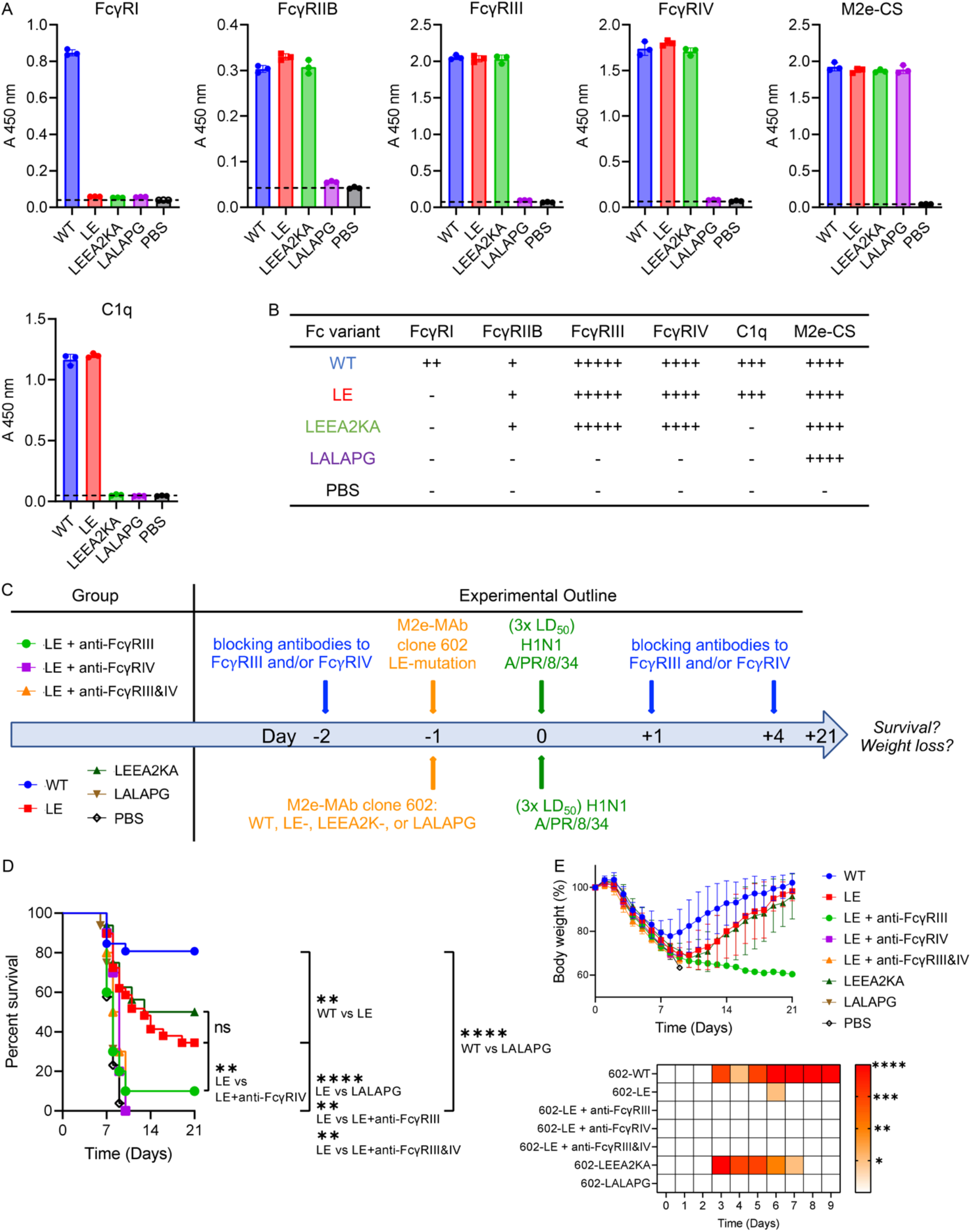
IgG2a M2e-MAbs mediate critical effector functions through FcγRI, III, and IV when administered systemically to influenza A virus-challenged mice. (**A**) Recombinant mouse FcγRI, FcγRIIB, FcγRIII, FcγRIV, or the M2e-consensus sequence (CS) peptide (5 µg/ml) was used as a coating antigen. WT M2e-MAb clone 602, and clone 602 with either the LE, LEEA2KA, or LALAPG Fc-domain mutation (50 µg/ml), were used to determine binding to recombinant FcRs or the M2e-CS peptide by direct ELISA. PBS was used as a control. N=3 independent experiments. To test the binding of the WT and Fc-mutant M2e-MAb clone 602 preparations to C1q, the WT or mutant antibodies were used as coating antigens (5 µg/ml), followed by mouse C1q protein (0.5 µg/ml). PBS was used as a control. Anti-mouse C1q-biotin and avidin-HRP were then used to quantify C1q protein bound by the capture antibodies by indirect ELISA. N=3 independent experiments. (**B**) Summary of the differential binding affinities of the 602-WT and Fc-variants to recombinant FcγRs, M2e-CS, and C1q as determined in (A). (**C**) Experimental outline: Groups of 6-8-week-old female Balb/c mice were prophylactically intraperitoneally injected with either the wild type (IgG2a) or the indicated Fc-mutant M2e-MAbs (all clone 602) on day -1 (100 µg/mouse). Control mice received PBS. Twenty-four hours later (on day 0), mice were infected with a lethal dose (3x LD50) of PR8 by aerosol inhalation. Additional groups of mice were intraperitoneally injected with blocking MAbs specific to FcγRIII (100 µg/mouse) and/or FcγRIV (200 µg/mouse) on days -2, 1, and 4 and additionally prophylactically treated with the M2e-MAb clone 602 with the LE mutation (100 µg/mouse i.p.) at day -1. Mice were challenged with a lethal dose (3x LD50) of PR8 by aerosol inhalation on day 0, and (**D**) survival and (**E**) weight loss was monitored for 21 days post-infection. Statistical significance for weight loss is shown in the heat map. N=10-29 mice per group; * p < 0.05, ** p < 0.01, *** p < 0.001, and **** p < 0.0001; Survival: Log-rank Mantel-Cox test; Weight loss: two-way ANOVA with Dunnett’s multiple comparison test.

### A single therapeutic treatment of mice with the triple M2e-MAb significantly ameliorates disease severity and enhances survival in mice challenged with H1N1 PR8 or with the pathogenic avian influenza strain A/Anhui/1/2013 (H7N9)

Last, we examined if treating mice with the 472/522/602 IgG2a triple cocktail enhances their survival when the therapy is administered after lethal H1N1 A/PR/8/34 infection (**Fig. 7**). We tested the therapeutic effect of our IgG2a-M2e-MAb triple cocktail therapy using two IAV challenge doses, one 100% lethal (**Fig. 7 A**), the other 75% lethal (**Fig. 7 B**). We established that the therapeutic treatment of Balb/c mice with the 472/522/602 IgG2a triple cocktail significantly enhanced the survival of Balb/c mice when administered on the day of infection, as well as one and two days later. At lower viral challenge doses (75% lethality), disease severity, as determined by weight loss, was significantly ameliorated when the M2e-MAb triple cocktail therapy was administered as late as day 3 after the IAV challenge. Survival was also improved in all therapeutically treated experimental groups (days 0-4), albeit statistical significance was not achieved for all time points.

**Figure 7.**
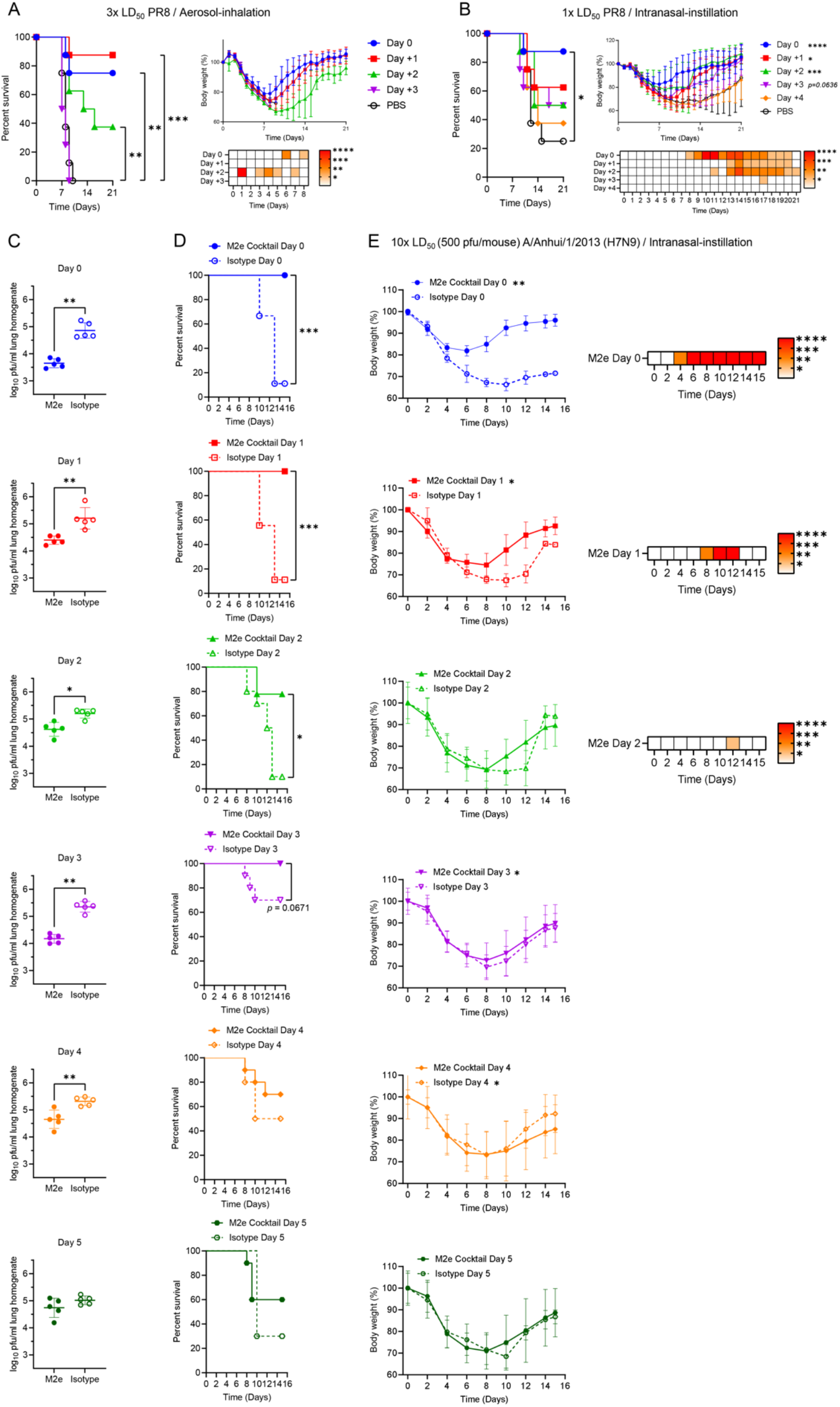
A single therapeutic treatment of mice with the triple M2e-MAb significantly ameliorates disease severity and enhances survival in mice challenged with H1N1 PR8 or with the pathogenic avian influenza strain A/Anhui/1/2013 (H7N9). (**A**) Groups of 6–8-week-old female Balb/c mice were infected with (**A**) a lethal dose (3x LD_50_) of PR8 by aerosol-inhalation or (**B**) a sub-lethal dose of (1x LD_50_) of PR8 by intranasal instillation. All mice received 450 µg in total of the M2e-MAb triple cocktail one-time at the indicated time point (150 µg each of clone 472, 522, and 602). Survival and weight loss were monitored for 21 days post-infection. A Mantel-Cox log rank test was used for the survival analysis. To compare percent weight loss, a one-way ANOVA (overall weight loss) or a two-way ANOVA (weight loss on individual days post-infection) with Dunnett’s multiple comparisons test was used to compare multiple experimental groups with one isotype control group. (**C**-**E**) Groups of 6–8-week-old female Balb/c mice were infected with a lethal dose (10x LD50) of H7N9 Anhui1 by intranasal instillation. All mice received either 450 µg of the M2e-MAbs triple cocktail or isotype control one-time at the indicated time point. (**C**) Survival and (**D**) weight loss were monitored for 15 days post-infection, and (**E**) lung viral titers were measured on day 5 (on day 6 for the Day 5 groups) by plaque assay. Statistical significance for weight loss is shown in the heat map. N=8-10 mice per group for survival and weight loss. N=5 mice per group for lung viral titers. A Mantel-Cox log rank test was used for the survival analysis. A paired t test (overall weight loss) or a two-way ANOVA with a Sidak’s multiple comparisons test (weight loss on individual days post-infection). Mann-Whitney test was utilized for lung viral titers on the specified day. * p < 0.05, ** p < 0.01, *** p < 0.001, and **** p < 0.0001.

To examine the M2e-MAb cocktail’s therapeutic efficacy to an IAV serotype of significant public health concern, we also evaluated the M2e-MAb cocktail’s therapeutic efficacy in mice challenged with the avian influenza virus H7N9, which causes severe disease and high mortality in poultry and humans ^6^. Importantly, treating H7N9-challenged Balb/c mice once with the M2e-Mab cocktail significantly lowered lung viral titers, even when the treatment was administered as late as day 4 p.i. (**Fig. 7 C**). The reduction in lung viral titers correlated with significantly improved survival: 100% of H7N9 challenged, and triple M2e-MAb cocktail treated Balb/c mice survived when the M2e-MAb treatment was administered on the day of infection (day 0), or day 1 or day 3 p.i. (**Fig. 7 D**). Also, 80% of mice treated on day 2, 70% of mice treated on day 4, and 60% of mice treated on day 5 p.i. survived (**Fig. 7 D**). It is noteworthy that statistical significance was not obtained at the later time points (days 3-5 p.i.) due to the improved survival of the isotype control groups, which benefitted from hydration when IgG2a-isotype control MAb was infused in 200 µl of saline on day 3, 4, or 5 post-H7N9 challenge. However, survival was improved and remained high in all M2e-MAb treated experimental groups (days 0-5 p.i.) compared to IgG2a isotype control MAb infusions. Disease severity, as determined by overall weight loss, was also significantly ameliorated when the M2e-MAb triple cocktail therapy was administered as late as day 4 after the H7N9 challenge (**Fig. 7 E, overall weights**). These data establish our triple M2e-MAb cocktail therapy as robustly effective against one of the most lethal IAV serotypes.

## DISCUSSION

Seasonal influenza virus infections burden the human population and kill about half a million people annually worldwide, including about 50,000 people in the US ^7^. Thereby, influenza virus infections cost the US economy an estimated 11.2 billion US dollars ^8^. Of the six FDA-approved treatments for IAV ^9, 10^, viral escape mutants have been found in response to each therapy ^11–14, 18, 19^, some already during clinical trials ^18^, with select mutations increasing the transmissibility of resistant viruses ^16^. For example, Amantadine, and Rimantadine, matrix protein 2 (M2) channel blockers, are ineffective in treating H1N1 and H3N2 infections due to widespread resistance ^11^. To better prepare for seasonal and pandemic IAV outbreaks, a safe, effective, and universally protective “off-the-shelf” treatment option is critically needed ^67^. The ectodomain of the IAV-encoded M2 protein is a suitable target for such a therapy: M2 assembles into a highly conserved proton channel expressed on influenza virions and infected cells, is required for infection and the viral life cycle, and its N-terminus is highly conserved across different IAV serotypes ^30–32^. However, previous mouse and human M2e-MAbs with protective potential are either strain-specific, have limited efficacy, or drive viral immune escape, and none have been successfully developed clinically ^36, 56^. To address this deficit, we used AuNP-M2e-CpG vaccination of Balb/c mice ^53^, a vaccine strategy resulting in life-long antibody-mediated immunity ^68^, to generate several M2e-specific MAb clones that are broadly protective against lethal IAV infection in mice ^53^.

In this study, we developed a highly and broadly effective viral escape mutant-resistant triple M2e-MAb cocktail therapy. We used rigorous molecular, virologic, and immunologic approaches to map each M2e-MAb triple cocktail component antibodies binding site in the highly conserved N-terminus of the M2 ectodomain and determined their competitive binding abilities. To enable protection at lower doses, increase universality, and resistance to viral immune escape, we tested double and triple M2e-MAb cocktails composed of individual M2e-MAb components that each protect mice from lethal IAV challenge. We demonstrated the broad therapeutic applicability, minimum effective dosage, and therapeutic administration time points for our M2e-MAb triple cocktail in mice challenged with human and zoonotic BSL-2 and BSL-3 IAV strains. To our knowledge, we are the first to have developed a universally effective and viral escape mutant-resistant M2e-MAb triple cocktail that significantly reduces lung viral titers and ameliorated disease severity even when administered as late as 4 days p.i. at least for the HPAI serotype H7N9.

Several considerations are noteworthy: First, while low-dose combinations of two M2e-MAb pairs failed to protect IAV-challenged mice from lethality completely, a triple cocktail comprised of M2e-MAbs with competitive binding sites in the M2-proteins N-terminal region was universally protective and highly effective at low doses. Thus, the protection provided by the triple M2e-MAb cocktail is not due to a dominant effect by one or two of the component antibodies but requires the contributions of all three M2e-MAbs cocktail component antibodies. The concept of combining multiple therapies or combining epitopes has been suggested previously ^25, 69^. However, to our knowledge, this theory had not yet been demonstrated for IAV. Thus, our study is the first to demonstrate a combination of MAbs for IAV treatment to be more effective than individual MAb therapies.

Second, using a triple cocktail may allow for lowering of the therapeutic dose with an additional benefit: When administered prophylactically, the 60-µg dose (approximately 3.7 mg/kg) is 100% protective against lethal PR8 challenge. In contrast, previously examined IAV-M2e-specific antibodies, such as TCN-032, were 60% protective when administered trice at 24.5 mg/kg on days 1, 3, and 5 post-IAV challenge ^37^. Estimates based on our data for a starting prophylactic dose for a human clinical trial would be 0.3 mg/kg ^70^, a dose over 130 times lower than the therapeutic dose used in the TCN-032 clinical trial ^37, 70^. Though, the efficacy and associated therapeutic dosage of our M2e-MAb triple cocktail remain to be determined in humans.

Third, our M2e-MAbs are not broadly neutralizing. We observed partial neutralization of H1N1 and H7N9 in prophylactically triple M2e-cocktail treated virally infected mice, presumably due to clone 472, which by itself has some neutralizing activity ^53^. However, the therapeutic effect of the M2e-MAb cocktail was most robust when expressed as the IgG2a isotype, and therapeutic efficacy depended on FcγRI, FcγRIII, and FcγRIV mediated effector functions. Of note, we used blocking antibodies to block FcγRIII, and FcγRIV functions, rather than genetically deficient animals, as FcγRIII-deficiency may lead to significant compensatory overexpression of FcγRIV on macrophages and circulating neutrophils in FcγRIII-deficient mice, as determined by real-time RT-PCR and flow cytometry ^42, 66^. In contrast, FcγRIV expression does not differ between FcγRI- deficient and control animals ^66^. While not as pronounced, FcγRIV-deficiency in mice also results in a slightly increased FcγRIII expression ^42^. While some M2e-specific antibody clones have neutralizing activity ^57, 62, 71^, those who do neutralize certain IAV strains do so at the cost of universality ^57, 62, 71–74^. Most previously described M2e-MAbs do not neutralize IAVs ^53, 63, 74–78^. Neutralization, while effective, is not a requirement for protection from lethal IAV challenge, as demonstrated here and by other non-neutralizing but protective antibodies to M2e-MAbs ^63, 78^ and influenza’s HA stalk region ^63, 79, 80^, which depend on a variety of FcR-mediated effector function for their protection mechanisms ^63, 81, 82^. While we identified FcγRI, FcγRIII, and FcγRIV as essential for IgG2a M2e-MAb mediated protection, we did not attempt to identify which FcR- functions are mediated by these receptors. FcγRI, FcγRIII, and FcγRIV are involved in phagocytosis, degranulation, and ADCC. NK cells, monocytes, macrophages, neutrophils, dendritic cells, basophils, mast cells, and eosinophils express FcγRIV, while FcγRIII is expressed by NK cells, monocytes, and macrophages, and FcγRI by monocytes, macrophages, and dendritic cells ^83, 84^. Due to the many potentially involved (combinations of) immune cell types, we did not attempt to identify them and their specific FcR-mediated effector functions in mice. While the selective depletion of some tissue-resident subsets of immune cells or their isolations and adoptive transfers have been done ^63^, markers that are exclusively expressed by a single immune cell type are rare. Thus, these approaches either fail to distinguish related and phenotypically similar immune cell types or result in too few cells for adoptive transfer. Also, as any M2e-MAb-based therapies in humans would utilize a humanized M2e-MAb cocktail where human isotypes interact with human FcRs, thus allowing the engineering of the MAb-based therapy to optimize antibody half-life ^85^ and FcR-mediated effector functions ^86^.

Most importantly, we demonstrated individual M2e-MAbs and the resulting triple cocktail as viral escape mutant resistant treatments in immunocompetent and immunodeficient mice, using both Sanger sequencing studies of the M-region and *in vivo* challenge studies demonstrating unchanged susceptibility of the virus to the M2e-MAb triple cocktail therapy after prolonged viral passage in the presence of therapy. Thus, despite IAV’s well-established ability to develop escape mutations to M2e-MAbs *in vitro* ^57^ and in mice ^57, 61–63^, our data demonstrate that it is possible to target IAVs with M2e-MAbs without driving viral immune escape and that we have generated such a product. Another crucial attribute of our triple M2e-MAb cocktail therapy is its ability to effectively reduce lung viral titers, ameliorated disease severity, and reduce lethality when administered up to 4 days p.i. to Balb/c mice challenged with the virulent H7N9 avian influenza virus. In contrast to our data, FDA-approved influenza virus therapies must be administered within 48 hours of symptom onset ^67^, and therapeutic efficacy has not been demonstrated for published M2e-specific MAbs administered later than 2 days p.i. ^37, 52^. Emerging, re-emerging, and smoldering outbreaks of zoonotic influenza viruses, including the H7N9 viruses pose a significant public health threat to the human population due to their ability to cause upper and lower respiratory tract disease, severe pneumonia with respiratory failure, encephalitis, and multi-organ failure ^6^. Thus, our results are impactful as they establish that robust and effective off-the-shelve MAb-based therapeutics can be developed to protect us from future potential influenza virus pandemics.

In summary, our study establishes a triple cocktail of cross-protective M2e-MAbs to be 1) efficacious at preventing IAV lethality at low doses, 2) consistently and universally protective and therapeutic between IAV strains, and 3) resistant to viral immune escape. These desirable attributes make the M2e-MAb cocktail a strong candidate for a universal “off-the-shelf” IAV therapeutic, ensuring rapid availability, consistent protection between strains, and prevention of viral resistance. Broadly effective, escape mutant-resistant off-the-shelf therapeutics may be our best option for reducing lethality during future influenza virus pandemics and may provide us with the necessary window to develop and disseminate a pandemic-serotype-specific vaccine. To our knowledge, our data are the first demonstration of a successful universally protective and viral escape mutant-resistant influenza A virus-specific M2e-MAb cocktail. Thereby, our data will critically shape future M2e-MAb-based influenza-therapeutic development.

## MATERIALS AND METHODS

### Animals and Approvals

Female 6-8 week old Balb/c and RAG2-KO mice were ordered from Charles River Laboratories Inc., San Diego, and Envigo RMS, Inc., Indianapolis, Indiana, USA. Mice were cared for in the animal facilities of the Texas Children’s Hospital, the Scripps Research Institute, and the University of Georgia, Athens. All protocols involving the use of experimental animals in this study were approved by Baylor College of Medicine’s, The Scripps Research Institute’s, and the University of Georgia, Athens’ Institutional Animal Care and Use Committees and were consistent with the National Institutes of Health Guide for the Care and Use of Laboratory Animals. All experiments using H7N9, or H5N1 avian influenza virus were reviewed and approved by the institutional biosafety program at the UGA and were conducted in biosafety level 3 enhanced containment. Work with highly pathogenic avian influenza virus H5N1 followed guidelines for using Select Agents approved by the CDC.

### Viruses

Influenza strains used in this study were A/Puerto Rico/8/1934 (H1N1; PR8), A/California/07/2009 (pdmH1N1; CA07), A/Vietnam/1203/2004 (H5N1; VN1203), and A/Anhui/1/2013 (H7N9; Anhui1). Before use in this study, all viruses were obtained, passaged, isolated, and quantified as previously described ^53^.

### Intranasal IAV challenge

PR8 was administered intranasally in 20 μl of PBS to mice anesthetized with isoflurane. CA07 and VN1203 were administered intranasally in 30 μl of PBS to mice anesthetized with Ketamine/xylazine. Anhui1 virus was administered intranasally to mice anesthetized with 2,2,2- tribromoethanol in tert-amyl alcohol (Avertin; Aldrich Chemical Co). Each challenge with CA07, VN1203, and Anhui1 viral inoculum was back-tittered on MDCK-ATL cells to confirm the dose. All animals were monitored for body weight and humane endpoints for euthanizing. Survival and weight loss were monitored for up to 21 days post-infection or until all animals recovered to at least 90% of the starting body weight.

### Aerosolized IAV challenge

Inside a BSL-2 safety cabinet, PR8 in 1X PBS and placed into a MiniHEART-HiFlo^®^ Continuous Nebulizer (SunMed). An Eisco^TM^ Superior Stand and Rod Set with a three-prong clamp with a Boss Head (Fisher Scientific) was used to stabilize the nebulizer. The nebulizer inlet was connected to a Flow Gauge and EasyAir2 compressor (Precision Medical) using oxygen tubing (SunMed). Corrugated plastic tubing (Harvard Bioscience) was attached to the top of the nebulizer and the other end to the mouse container. The in-unit flow gauge of the compressor was set to 10 L/min, and the flow gauge, attached to the condenser, to 8 L/min. Mice were exposed to PR8- containing aerosol vapor for 25 minutes for infection with the aerosolized PR8 virus. Overflow influenza aerosol was disposed of into a waste container containing 10% bleach through the oxygen tubing.

### Viral titer measurement

A subset of IAV-infected mice was humanely euthanized, and tissues were collected for virus titer 3 days post-infection. Lung tissue samples were tittered by plaque assay. Supernatants were serially diluted 10-fold in DMEM and added to MDCK.2 cells (ATCC) in 6 or 12-well plates tissue culture plates. After a 1 to 2-hour culture, 2 ml of a 0.27% agar in DMEM containing 0.5 µg/ml of TPCK-trypsin or 1.2% Avicel microcrystalline cellulose overlay (MEM supplemented with HEPES, L-Glutamine, NaHCO3, Penicillin/Streptomycin/Amphotericin B, and 2 µg/ml of TPCK-trypsin) was overlaid. Plates were incubated for 48 to 72 hours at 37℃ with 5% CO2, washed, fixed with 4% paraformaldehyde or methanol:acetone (80:20), and stained with 0.4% crystal violet solution to visualize plaques. For H5 or H7 viruses, supernatants were diluted in DMEM+2% FBS, and Avicel overlay included 2% FBS instead of TPCK trypsin.

### Cell lines

FreeStyle^TM^ 293-F cells were purchased from ThermoFisher Scientific. FreeStyle^TM^ 293 Expression Medium (ThermoFisher Scientific) was used for culturing and transfecting the cells without any additional reagents in orbital shakers at 135 rpm, 37℃, and 8% CO2. MDCK.2 cells were purchased from ATCC and cultured in DMEM containing 10% FBS, 25 mM HEPES, 4 mM L-Glutamine, and 100 U/ml of Penicillin-Streptomycin. Consensus and Vietnam matrix protein 2 (M2)-inducible HEK cells were generated using the Flp-In™ T-REx™ 293 Cell Line system (Invitrogen) and grown following the methods in the previous papers ^53, 87^. The HEK cells were cultured in DMEM containing 10% FBS, 25 mM HEPES, 4 mM L-Glutamine, 100 µg/ml of Hygromycin B, 15 µg/ml of Blasticidin, and 100 U/ml of Penicillin-Streptomycin. Both MDCK.2 cells and the HEK cells were maintained at 37℃ and 5% CO2.

### Neutralization assay

MDCK.2 cells (ATCC) were seeded at a density of 1 × 10^6^ cells/well in 6-well plates (VWR International) and incubated overnight at 37℃ 5% CO2. 20 – 50 pfu/ml PR8 was incubated with 25 µg/ml of the indicated single M2e-MAb or the triple M2e-MAb cocktail (8.3 µg/ml of each cocktail component antibody) at 4℃ for 30 min in PBS. MDCK.2 cells were washed twice with PBS before antibody-virus mixtures were added, and the plates were gently shaken every 10 – 15 min at 37℃ for 1 h. Then, cells were overlayed with 0.27% agar in DMEM containing 0.5 µg/ml of TPCK-trypsin and incubated at 37℃ for 3 days before plates were fixed with 4% paraformaldehyde for 1 hour and stained with 0.4% crystal violet solution for 30 min. Clear plaque numbers were counted, and virus titers (Log10 pfu/ml) were calculated.

### M2e-MAbs production from hybridomas

Antibody production for 472, 522, and 602 clones was performed by expanding the hybridomas as previously described ^53^. IgG2a isotype control-matched antibody was purchased from BioXCell (#BE0085).

### Biotinylation of M2e-MAbs

M2e-MAbs were biotinylated using EZ-Link Hydrazide Biocytin (ThermoFisher Scientific) according to the manufacturer’s instructions for labeling glycoproteins with hydrazide biocytin. Biotinylated M2e-MAb was separated from non-reacted material by dialysis (10kd MWCO; ThermoFisher Scientific) in 1X phosphate-buffered saline (PBS) for 12 hours. Samples were removed from dialysis cassettes, aliquoted, and stored at 4°C.

### Cloning, expression, and purification of isotype-switched M2e-MAbs

Variable region sequences of the M2e-MAbs were verified by Sanger sequencing. For the M2e-MAbs heavy chain and light chain variable regions, gBlock double-stranded DNA was synthesized and purchased from Integrated DNA Technologies (IDT, USA). AgeI-HF/Eco47III (NEB) or AgeI-HF/BstAPI (NEB) enzyme sites were added at the end of the gBlock DNA for their cloning into mouse IgG1 or IgG2a heavy chain plasmids, or the mouse kappa light chain plasmid (InVivoGen). The gBlock DNA and plasmids were cut with the specified enzymes, subjected to agarose-gel electrophoresis, extracted using the Monarch^®^ DNA Gel Extraction Kit (NEB), ligated using T4 DNA Ligase (Promega), transformed into NEB^®^ 10-beta Competent *E. coli* cells (NEB), and colonies screened with Zeocin for the heavy and light chain plasmids using Blasticidin (InVivoGen). The plasmids were purified using QIAprep Spin Miniprep Kit (Qiagen), and their sequences were confirmed by Sanger sequencing (Genewiz). Log phase (0.3 – 3 × 10^6^ cells/ml) FreeStyle^TM^ 293-F cells were co-transfected with the specified M2e-MAbs heavy and light chain variable region plasmids (InvivoGen) using Polyethylenimine (PEI, Polysciences). After 5 days, cells were centrifuged, and antibodies purified from the supernatants using Protein G or A bead affinity chromatography by The Scripps Research Institute’s Antibody Core Facility.

### Epitope mapping using M2e-peptide libraries

M2e peptide libraries were synthesized based on the M2e-consensus sequences (CS) peptide, N- SLLTEVETPIRNEWGCRCNDSSD, and purchased from Genscript. The specified M2e peptide library was coated in the 96-well plates (Corning) and incubated in 15 mM Na2CO3 and 35 mM NaHCO3 bicarbonate buffer (pH 9.6) at 4℃ overnight. The plates were blocked with PBS containing 1% BSA for 1 hour at room temperature before the specified biotinylated IgG1 M2e-MAbs or biotinylated IgG1 control MAbs (Clone: MOPC-21, BioXCell) diluted in PBS 0.1% Tween 20 (PBS-T) were added and incubated for 1 hour at room temperature. The plates were washed 3 times with PBS-T, and Streptavidin-HRP (Pierce Chemical), diluted in PBS-T, was added to the plates. After 1 hour, the plates were washed 4 times with PBS-T, and the peroxidase substrate tetramethylbenzidine TMB solution (Fisher Scientific) was added. After 5 minutes, 2N H2SO4 was added in the plates and absorbance detected at 450 nm using the SpectraMAX^®^ iD3 instrument (Molecular Devices).

### M2e-MAb competition ELISA

Influenza strains used in this study were A/Puerto Rico/8/1934 (H1N1; PR8), A/California/07/2009 (pdmH1N1; CA07), A/Vietnam/1203/2004 (H5N1; VN1203), and A/Anhui/1/2013 (H7N9; Anhui1). Before use in this study, all viruses were obtained, passaged, isolated, and quantified as previously described ^53^. Nunc Maxisorp Flat-Bottom plates (ThermoFisher Scientific) were coated overnight at 4°C with the indicated purified inactivated influenza virus at 0.5 μg/ml in bicarbonate buffer (pH 9.6). After washing the plates 3 times with PBS 0.05% Tween 20, the plates were blocked with 1% BSA in PBS for 2 hours and washed 3 times with PBS 0.05% Tween 20 before biotinylated M2e-MAbs were added at 2 μg/ml (the concentration resulting in approximately 50% saturation in the assay) to the plates and incubated for 1 hour at 37°C. Then, the indicated M2e-MAbs were added as the competing antibody in 4- fold dilutions and incubated for 1 hour at 37°C. After washing the plates with PBS 0.05% Tween 20 3 times, a 1:10,000 Streptavidin-HRP dilution (Vector Laboratories) was added to the plates and incubated for 1 hour at 37°C. The plates were then washed with PBS 0.05% Tween 20 3 times, and TMB substrate was added for 10 – 15 min before the reaction was stopped with the addition of H2SO4 and absorbance measured at OD 480 nm.

### M2e-MAb analysis by SDS-PAGE and Coomassie staining

SDS-PAGE of the indicated M2e-MAbs prepared in both reducing and non-reducing conditions was performed using NuPAGE^TM^ Bis-Tris Welcome Pack, 4-12%, 10-well (Fisher Scientific). Protein size was compared to PageRuler™ Plus Prestained Protein Ladder (10 to 250 kDa). To visualize proteins, the SDS-PAGE gel was stained with Coomassie Stain Solution (Rockland) for 30 min, and then de-stained with Coomassie Brilliant Blue R-250 Destaining Solution (Bio-Rad), and bands visualized using a ChemiDoc XRS+ System (Bio-Rad).

### Detection of M2e-MAb binding to M2-expressing HEK cells by flow cytometry

M2 expressing HEK cells were treated with 2 µg/ml of tetracycline to induce M2 expression. After 48 h, the cells were detached with Trypsin-EDTA (0.05%), washed with PBS containing 2% FBS, and stained with the specified antibodies for 30 min at room temperature. After washing with PBS containing 2% FBS, the cells were stained with Alexa Fluor 488-conjugated goat anti-mouse IgG secondary antibody (Invitrogen) for 30 min, washed, and fixed in 2% paraformaldehyde. Flow cytometry was performed to measure M2e-MAbs binding to M2-expressing HEK cells using Cytek^®^ Aurora (Cytek Biosciences).

### The 602-IgG2a Fc variants development

The pFUSE-CHIg-mG2a plasmid (InvivoGen) containing the 602 heavy chain variable region was used for the generation of Fc variant antibodies. gBlock double-strand DNAs for the L235E (LE), L235E/E318A/K320A/K322A (LEEA2KA), and L234A/L235A/P329G (LALAPG) mutated CH2 domains were synthesized and purchased from IDT with BamHI/BsrGI (NEB) restriction enzyme sites at the ends. The region encoding for the 602-IgG2a wild-type (WT) CH2 domain was replaced with the LE, LEEA2KA, or LALAPG gBlock DNA, all plasmids confirmed by Sanger sequencing, and each of the 602 Fc variants was produced and purified following the “Cloning, expression, and purification of isotype-switched M2e-MAbs” method section in the paper.

### Quantification of M2e-MAb binding to FcγRs, C1q, and M2e peptides by ELISA

96-well plates (Corning) were coated at 4℃ overnight with the indicated antigens, such as M2e peptides (M2e vaccine or consensus sequence), or recombinant mouse Fc gamma RI, RIIB, RIII, or RIV (R&D Systems) in 15 mM Na2CO3 and 35 mM NaHCO3 bicarbonate buffer (pH 9.6). Then, the plates were blocked with PBS containing 1% BSA for 1 hour at room temperature before the specified M2e antibodies, mouse IgG1 isotype control (Clone: MOPC-21, BioXCell), or mouse IgG2a isotype control (Clone: C1.18.4, BioXCell) were added in PBS-T (0.1% Tween 20) and incubated for 1.5 hours. Afterward, the plates were washed 3 times with PBS-T, and goat anti-mouse IgG-HRP (Jackson ImmunoResearch, USA) was added and incubated for 1 hour. Peroxidase activity was measured with TMB solution (Fisher Scientific), which was added for 5 minutes before 2N H2SO4 was added to stop the reaction. Absorbance was detected at 450 nm using a SpectraMAX^®^ iD3 instrument (Molecular Devices).

To perform the indirect ELISA to determine the binding of the M2e-MAbs to mouse C1q protein, 96-well plates were coated with the specified M2e antibodies in the bicarbonate buffer (pH 9.6) at 4℃ overnight. The plates were blocked with PBS containing 1% BSA. After 1 hour, the plates were washed 3-time with PBS-T, and added with mouse C1q protein (Complement Technology) in PBS-T. After a 1-hour incubation at room temperature, the plates were washed 3-time with PBS- T, added with biotinylated anti-mouse C1q antibody (Fisher Scientific), and incubated for 1 hour. Then, the plates were washed 3 times with PBS-T, added with HRP-conjugated streptavidin (Pierce Chemical), and incubated for 1 hour. The Plates were washed 4-time with PBS-T after the incubation with HRP-conjugated antibody or streptavidin, and peroxidase activity was measured with TMB solution (Fisher Scientific), which was added for 5 minutes before 2N H2SO4 was added to stop the reaction. Absorbance was detected at 450 nm using a SpectraMAX^®^ iD3 instrument (Molecular Devices).

### Prophylactic M2e-MAb treatment

As previously described ^53^, 24 hours before infection of mice with either a 5X or 10X LD50 of the specified IAV serotype, mice were prophylactically treated by intraperitoneal (i.p.) injection with the specified dose of the specified single M2e-MAb or the triple M2e-MAb cocktail.

### Triple M2e-MAbs cocktail treatment of influenza-infected Balb/c mice

Balb/c mice were challenged with 3X LD50 PR8 by aerosol-inhalation, or, alternatively, 1X LD50 PR8 by intranasal instillation, and 450 µg the triple M2e-MAbs cocktail was administered once, intraperitoneally, at the indicated time points after viral challenge. Survival and weight loss were monitored over 21 days post-infection. Balb/c mice were challenged with 10X LD50 Anhui1 by intranasal instillation, and 450 µg the triple M2e-MAbs cocktail or isotype control was administered in mice once intraperitoneally at the indicated time points after viral challenge. Survival and weight loss were monitored over 15 days post-infection. Lung viral titers were measured on day 5 post-infection (on day 6 for the Day 5 groups) by plaque assay.

### Viral M2e-MAb therapy escape mutant resistance assay and M-region sequencing

Twenty-four hours before IAV infection (day -1), groups of Balb/c mice were injected intraperitoneally with 60 µg of the indicated M2e-MAbs or the triple M2e-MAb cocktail (20 µg/each, 60 µg in total). On day 0, mice were anesthetized with isoflurane and intranasally infected with a 5X LD50 dose of PR8. On days three or four post-infection, mice were euthanized with an isoflurane overdose, and lungs were harvested to isolate the virus. IAV was isolated and purified from lung homogenates as previously described ^53^: In brief, lungs were placed in PBS on ice (0.75 ml/lung), homogenized, and centrifuged at 850g for 10 minutes at 4°C, after which the supernatant was placed in an Ultracel-100 tube (Amicon Ultra-15 centrifugal filter unit; Ultracel-100 regenerated cellulose membrane) and centrifuged at 4,000 g for 30 minutes at 4°C. The PR8- containing solution (remaining in the top of the filter tube) was placed in a new 15-ml tube and centrifuged at 850g for 5 minutes at 4°C. Then, 20 µl of the purified virus preparation was used to intranasally infect naïve Balb/c mice that had been treated with the indicated M2e-MAbs or the triple M2e-MAb cocktail 24 hours before infection. Alternatively, RAG2-KO mice were injected intraperitoneally with 60 µg of the indicated M2e-MAbs or the triple M2e-MAb cocktail (20 µg/each, 60 µg in total) 24 hours before and three days after intranasal infection with a 5XLD50 dose of PR8. Mice were sacrificed with an isoflurane overdose on day 7, lungs were harvested, and virus isolated as described above. PR8 was passaged six times through groups of seven Balb/c mice, or twice through groups of three Balb/c RAG2-KO mice before its isolation for Sanger sequencing. Total RNA was isolated using the QIAamp Viral RNA Mini Kit (Qiagen), cDNA synthesized using the M2-2 primer (5’-GCGAAAGCAGGTAGATATTG-3’), which binds to a 3’ noncoding region of influenza’s viral RNA segment 7 (vRNA7), and the Omniscript RT Kit (Qiagen). The cDNA was amplified by PCR using KAPA HiFi HotStart ReadyMix (Roche), using M2-2 and SEQ7 (5’-ATATCGTCTCGTATTAGTAGAAACAAGGTAG-3’) primers. The SEQ7 primer binds to a 5’ noncoding region of influenza vRNA7. The expected size of the PCR product is 1,042 bp, and this was confirmed by Gel electrophoresis. Genewiz, Inc performed sequencing analysis with M2SeqN1 (5’-ATGTTATCTCCCTCTTGAGC-3’) and SEQ7 primers. M2SeqN1 anneals to 331-351 of M1 cDNA, and SEQ7 anneals to a 5’ noncoding region of the cDNA. To identify potential viral escape mutants, the sequence of the “passaged” viruses were compared to the input PR8 virus (day 0), which matched the original M2e sequences ^88^.

### Identification of M2e-MAb-mediated Fc-effector functions required for the protection of PR8-challenged mice

To block the activities of FcγRIII and FcγRIV *in vivo*, Balb/c mice were intraperitoneally infused on days -2, 1, and 4 post PR8 infection with commercially available anti-FcγRIII antibody (100 µg/mouse; clone: 275003, R&D systems) and/or anti-FcγRIV antibody (200 µg/mouse; clone: 9E9 Biolegend), based on a published protocol ^89^. Indicated groups of mice were also therapeutically treated with a single intraperitoneal injection of either the 602-Fc-WT, LE-, or LEEA2K, or LALAPG M2e-Mabs (100 µg/mouse), or PBS as a control, on day -1 post PR8 infection, which was performed using a dose of 3X LD50 PR8 by aerosol-inhalation on day 0.

### Statistics

GraphPad Prism 9 was used for all statistical analyses. A Mantel-Cox log rank test was used for the survival analysis. To compare percent weight loss, a one-way ANOVA (overall weight loss) or a two-way ANOVA (weight loss on individual days post-infection) with Dunnett’s multiple comparisons test was used to compare multiple experimental groups with one isotype control group. A paired t test (overall weight loss) or a two-way ANOVA with a Sidak’s multiple comparisons test (weight loss on individual days post-infection) was used to compare a single experimental group with its corresponding isotype control group. A one-way ANOVA with Turkey’s multiple comparisons (comparing multiple experimental groups to a shared isotype control group) or Mann-Whitney test (comparing a single experimental group to its isotype control group) was utilized for lung viral titers on the specified day. All statistics are indicated in the figure legends. **** p<0.0001, *** p<0.001, ** p<0.01, * p<0.05.

## Supplemental Material

**Fig. S1:** M2e-specific antibodies bind to M2e competitively.

**Fig. S2:** M2e-MAb triple cocktail therapy reduces viral fitness and does not drive the development of viral escape mutants.

**Fig. S3:** Immunocompromised Rag2-KO mice significantly benefit from single M2e-MAb clone, alternating M2e-MAb clone, or M2e-MAb triple cocktail treatments.

**Fig. S4:** Isotype-switching, quality control, and functional analyses of isotype-switched M2e-MAb clones 472, 522, and 602.

**Fig. S5:** M2e-MAb therapy equally protects Balb/c mice infected with PR8 either intranasally or by aerosol inhalation.

**Table S1:** M2e-Consensus sequence (CS) alanine scanning peptide library

**Table S2:** M2e sequences of influenza A viruses isolated from WT or RAG2-KO mice after single M2e-MAb or M2e-MAb cocktail antibody treatments.

### Study approval

All institutions, each institutions’ Animal Care and Use Committees approved all protocols for animal experiments, and all institutions follow the "Principles of Laboratory Animal Care" formulated by the National Society for Medical Research.

## ACKNOWLEDGEMENTS

Funding for this work was supported by the National Institutes of Health [R01AI130065, 2017- 2021]; The Albert and Margaret Alkek Foundation, Houston TX [2015]; and the National Institute of General Medical Sciences of the National Institutes of Health under [Award Number AI053831]. This project was also supported by the Protein and Monoclonal Antibody Production Shared Resource at Baylor College of Medicine with funding from NIH Cancer Center Support Grant P30 CA125123. We thank Diane Kubitz and acknowledge the Center for Antibody Development and Production at The Scripps Research Institute for producing hybridomas and monoclonal antibodies used in this study. We thank Suzanne Epstein (Center for Biologics Evaluation and Research, U.S Food and Drug Administration, Silver Spring, MD) for providing A/PR/8/1934 (H1N1) and A/FM/1/1947- MA (H1N1). We also thank Earl G. Brown (University of Ottawa, Ottawa, Canada) for approving the sharing of A/FM/1/1947- MA (H1N1). We thank Ted Ross (University of Georgia, Athens, GA) for providing A/CA/07/2009 (H1N1) and Richard Webby (St. Jude Children’s Research Hospital, Memphis, TN) for providing A/Anhui/1/2013 (H7N9) and A/Vietnam/1203/2004 (H5N1). And we thank Jon Yewdell (National Institutes of Health, Bethesda, MD) for sharing the NP-specific hybridoma H16-L10. A/Anhui/1/2013 (H7N9) was provided via the WHO Global Influenza Surveillance and Response System (GISRS).

## AUTHOR CONTRIBUTIONS

SP and SMT conceptualized the project. SP, SMT, LB, and TK developed methodologies and designed experiments. LB, SLR, TK, AYS, SKJ, and CAJ performed experiments. SP, SMT, LB, SLR, and TK, AYS, SKJ, and CAJ analyzed data. LB and TK visualized data. SP and SMT acquired funding for the study and administered and supervised the project in their respective laboratories. SP, LB, and TK wrote the manuscript. SP, LB, TK, and SMT edited the manuscript.

---

**Figure S1.**
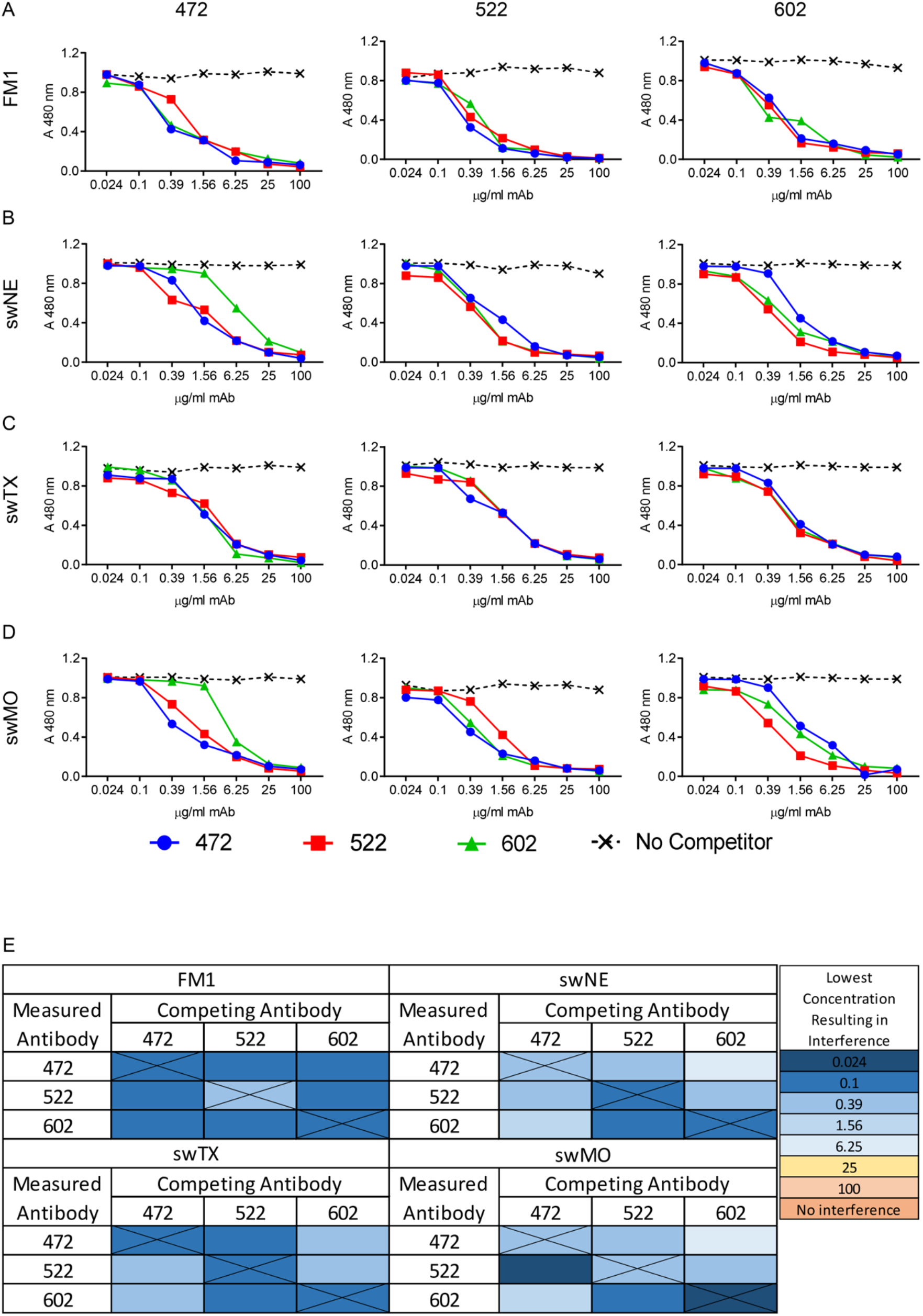
M2e-specific antibodies bind to M2e competitively. (**A-D**) Inactivated (**A**) FM1, (**B**) swNE, (**C**) swTX, or (**D**) swMO virions were used as the coating antigen for competition ELISAs. The competitive antibody was added at 4-fold dilutions starting with 100 μg/ml. The to-be-analyzed clone (clones 472 (IgG2a), 522 (IgG1), and 602 (IgG2a)) was biotinylated and added to the wells at a standard concentration of 2 μg/ml. Absorbance was measured using a biotin-specific secondary antibody. (**E**) Data summary: Shown are the concentrations for each antibody and virus at which the absorbance drops 0.1 absorbance units below the average absorbance of the “no competitor” control.

**Figure S2.**
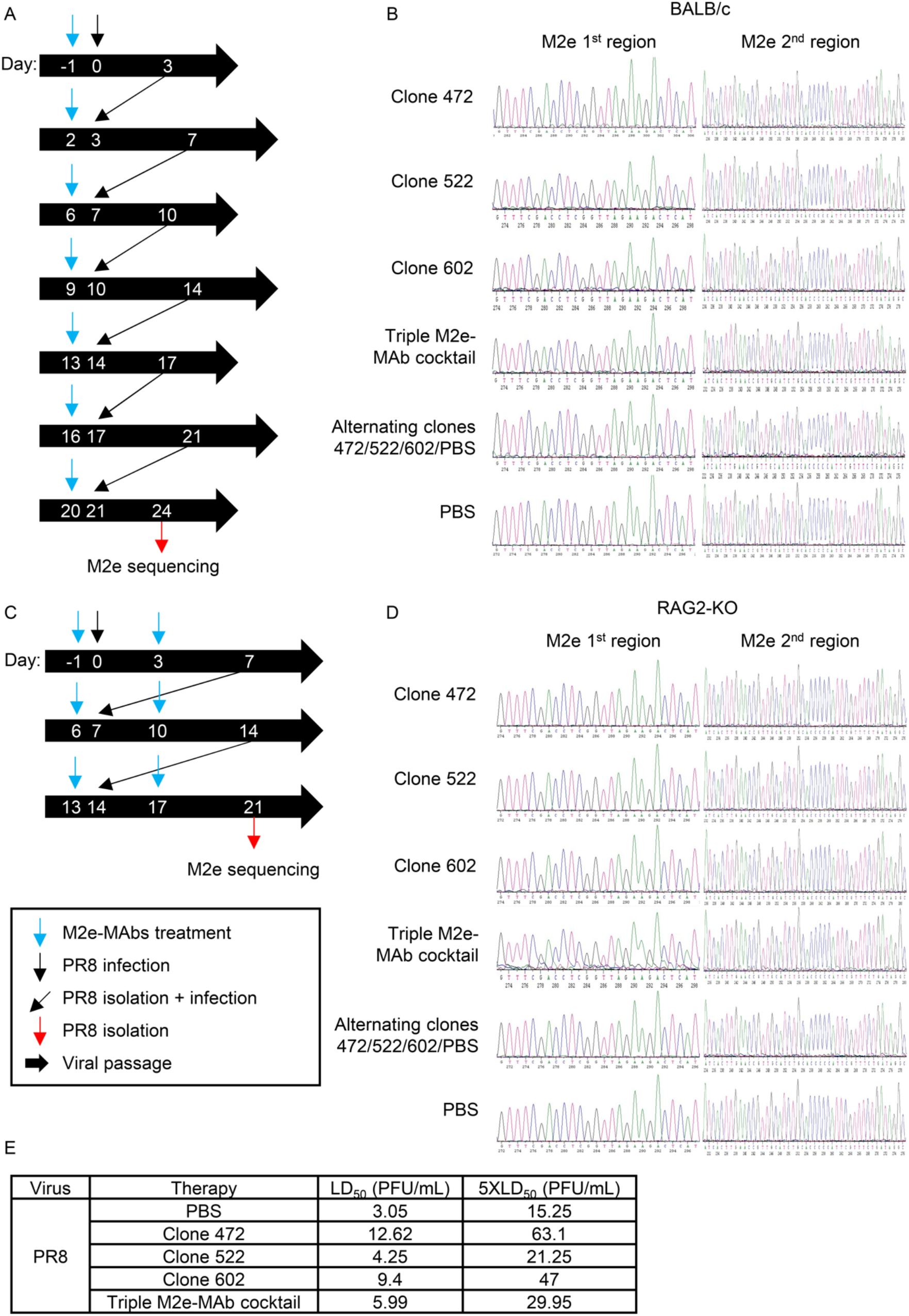
M2e-MAb triple cocktail therapy reduces viral fitness and does not drive the development of viral escape mutants. PR8 “stock virus” was passaged through WT or Rag2-KO mice, and the final viral isolates were analyzed by Sanger sequencing. (**A, C**) Outline of the time points for viral challenges of WT and RAG2-KO mice with virus isolated from control (PBS) or therapeutically treated groups of animals. PR8 stock virus was passaged (**A**) seven times (every 3- 4 days) through groups of Balb/c WT mice or (**C**) three times (every seven days) through groups of RAG2-KO mice for 24 or 21 days, respectively. Four mice per group were infected for each passage, except for the final passage, which used eight mice per group for both genotypes. At each passage, virus was isolated from lung homogenates of M2e-MAb triple cocktail (clones 472 (IgG2a), 522 (IgG1), and 602 (IgG2a)) or PBS control-treated mice and used to infect a group of naïve prophylactically M2e-MAb triple cocktail therapy or PBS (control) treated WT or Rag2-KO mice, as indicated. Intraperitoneal (IP) injections of M2e-MAbs treatments (60 µg total) of single MAbs (60 µg of clone 472 (IgG2a), 522 (IgG1), or 602 (IgG2a)), the triple M2e-MAb cocktail (20 µg each of clones 472, 522, and 602 = 60 µg total; 472-IgG2a, 522-IgG1, and 602-IgG2a), or alternating treatments with 472, 522, 602, or PBS (weekly in this order, 60 µg), or PBS control treatments are indicated by a blue arrow. (**B, D**) Sanger sequencing chromatograms of the indicated viral isolates. Sanger sequencing results for all experimental and control groups are reported in **Table S2.** (**E**) LD_50_ based on the viral challenge of Balb/c mice with 3 to 4 doses ranging from 0.24 to 30 PFU with the specified post-therapeutic passage PR8 viral isolates, as calculated using the Reed-Muench method ^90^.

**Figure S3.**
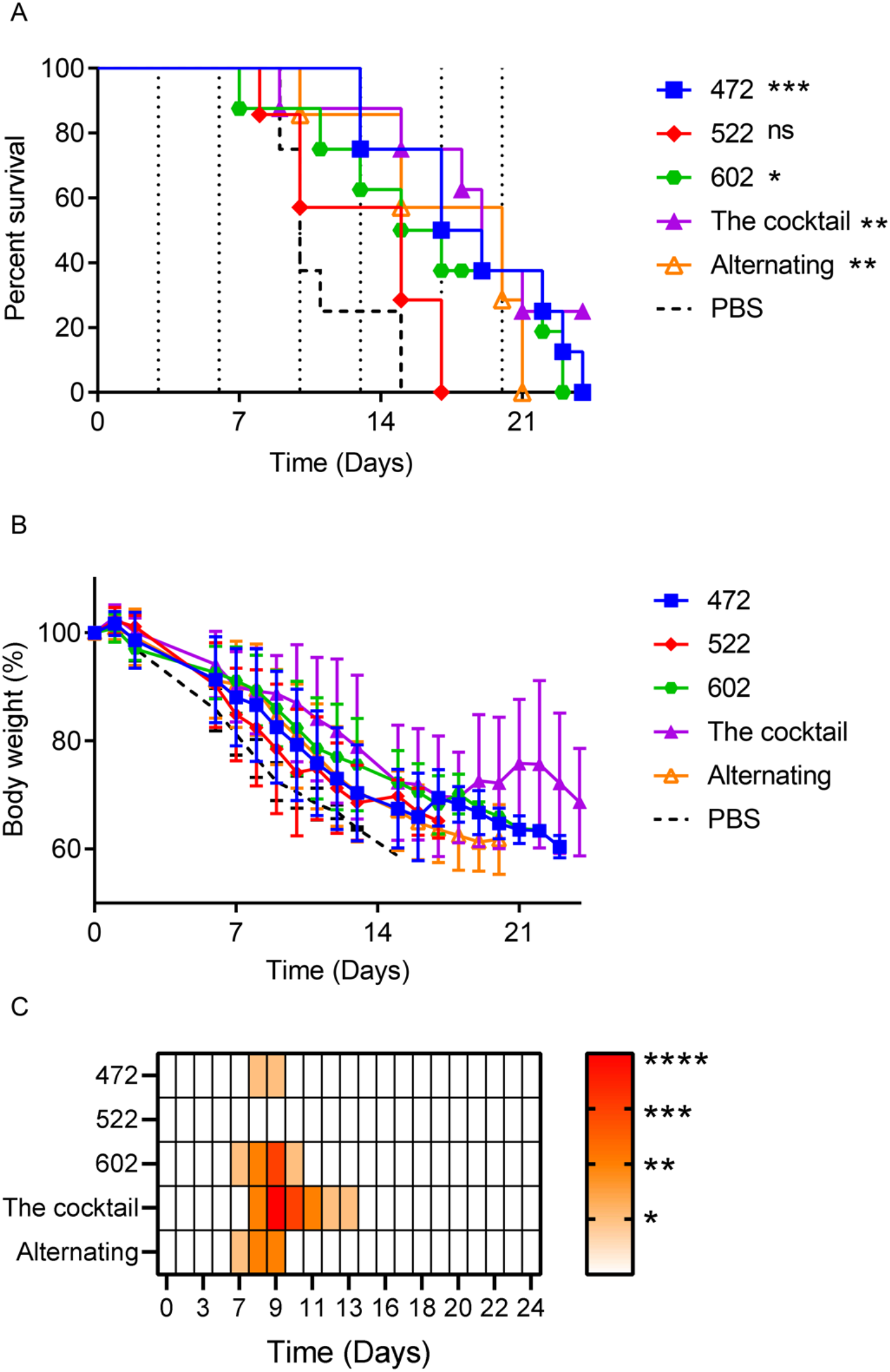
Immunocompromised Rag2-KO mice significantly benefit from single M2e-MAb clones, alternating M2e-MAb clones, or M2e-MAb triple cocktail treatments. (**A-C**) RAG2- KO mice were treated with a 60 μg dose of the indicated M2e-MAb clone (clones 472 (IgG2a), 522 (IgG1), or 602 (IgG2a)) or the M2e-MAb triple cocktail (clones 472 (IgG2a), 522 (IgG1), and 602 (IgG2a)) one day before infection with a sub-lethal dose (1x LD_50_) of PR8. Additional 60 μg treatments were administered twice weekly, alternating 3 and 4 days apart. Vertical dotted lines indicate treatments administered post-infection. For the alternating treatment, clones 472, 522, 602, or PBS were administered individually every X days, in this order, before alternating treatments were repeated. (**A**) Survival and (**B**) percent weight were determined for 24 days post-infection. (**C**) Heatmap indicating significant differences in the recorded percent weight loss as compared daily to the isotype control group. N=7-8 animals/group. Survival: Log-rank (Mantel-Cox) test; percent weight loss: Two-way ANOVA (Dunnett’s multiple comparisons) test. ** p<0.005, * p<0.05.

**Figure S4.**
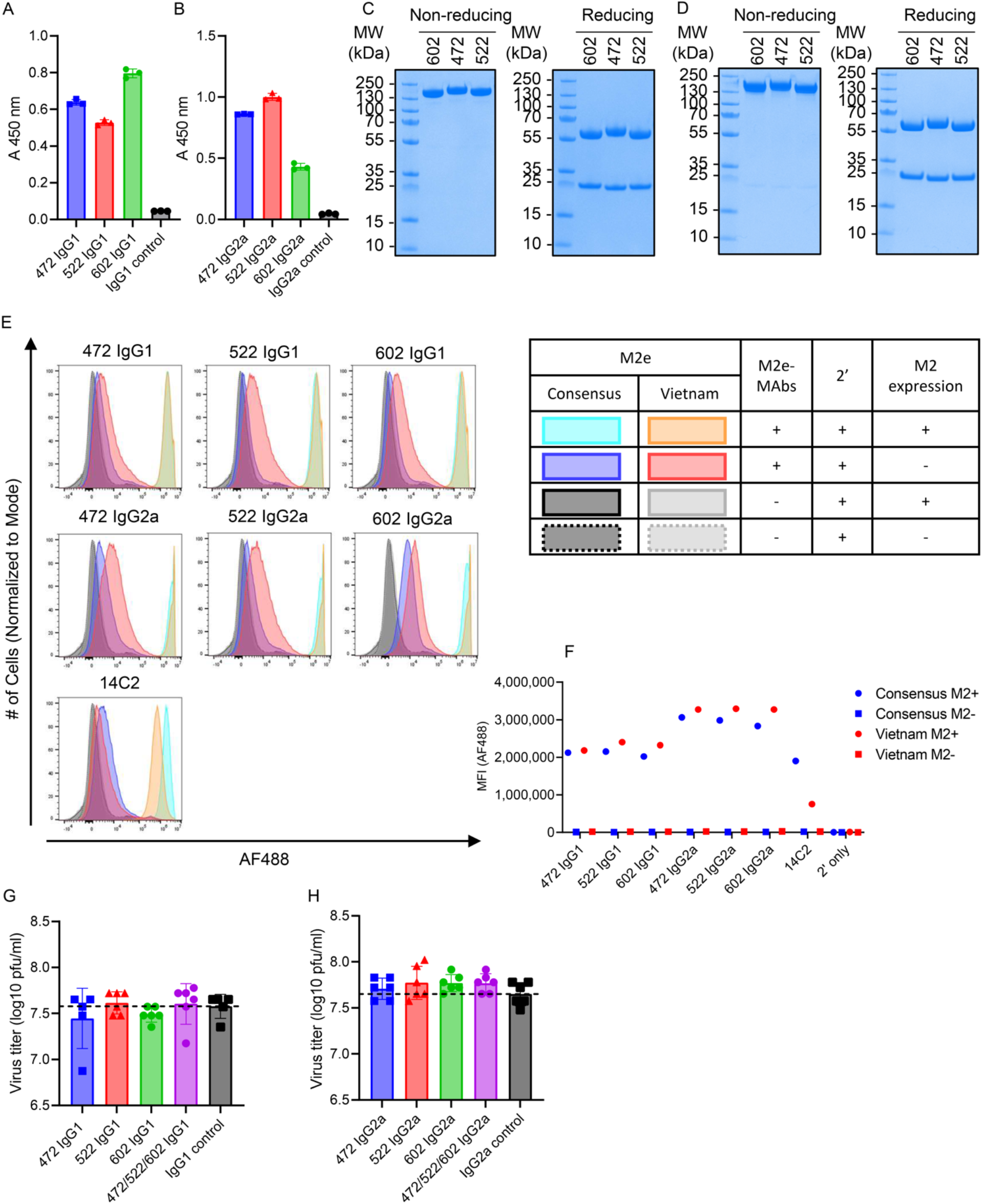
Isotype-switching, quality control, and functional analyses of isotype-switched M2e-MAb clones 472, 522, and 602. Clones 472, 522, and 602 were generated as IgG1 and IgG2a isotypes using the pFUSE2-CLIg-mk, pFUSE-CHIg-mG2a, and/or pFUSE-CHIg-mG1 expression vectors and the 293F expression system. (**A, B**) The binding of the isotype switched M2e-MAbs (2.5 µg/ml) to the M2e-vaccine sequences (VS) peptide (2.5 µg/ml) was determined by ELISA. IgG1 and IgG2a controls were used as negative controls. N=3 independent experiments. (**C, D**) Monomeric (**C**) M2e-MAbs IgG1 and (**D**) M2e-MAbs IgG2a (10 µg/antibody) were visualized by Coomassie staining in non-reducing and reducing conditions to confirm the correct size of their heavy and light chains. (**E**, **F**) The recognition of M2 on M2-CS and M2-Vietnam expressing HEK cells by the specified M2e-MAb clone or the commercially available M2e-specific MAb clone 14C2 (positive control) was determined by flow cytometry. Alexa Fluor 488-conjugated goat anti-mouse IgG was used as a secondary antibody to visualize M2e-MAb binding to M2-expressing HEK cells. (**E**) histogram overlay of experimental and control staining and (**F**) Alexa-488 mean fluorescent intensity (MFI). (**G**, **H**) The neutralization activity of the indicated M2e-MAb clones against the PR8 IAV serotype was determined by MDCK cell-based plaque assay. The specified M2e-MAb (single or cocktail) or isotype control (25 μg/ml) was incubated with PR8 (50 pfu/well) for 30 min at 4℃. MDCK cells were infected with the antibody-virus mixtures, cultured for 72 hours, stained with crystal violet, and plaque numbers counted. N=5-6 wells.

**Figure S5.**
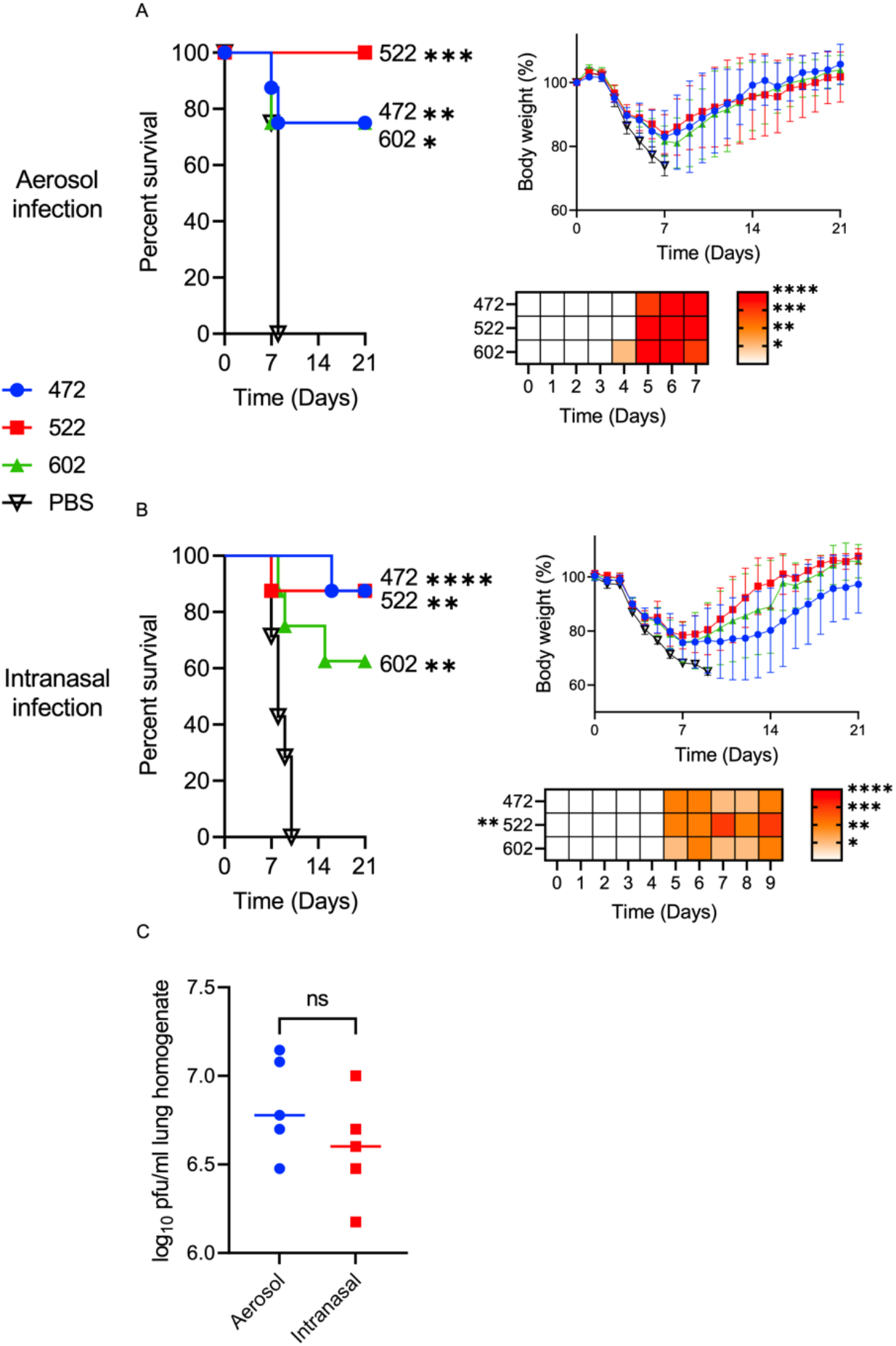
M2e-MAb therapy protects Balb/c mice infected with PR8 intranasally or by aerosol inhalation equally. Six to eight-week-old female Balb/c mice were prophylactically intraperitoneally infused with 100 µg of the M2e-MAb clone 472, 522, or 602 (all IgG2a isotype) or mock treated with PBS. One day later, one-half of the mice were infected by (**A**) aerosol-inhalation with 3x LD_50,_ the other half by (**B**) intranasal droplet challenge with 5x LD_50_ PR8, and their survival and weight loss were monitored for 21 days. N=7-8 mice per group. Survival: Log-rank (Mantel-Cox) test; weight loss: One-or two-way ANOVA (Dunnett’s multiple comparisons) test. Statistical significance for daily weight loss, as compared to the PBS control group, is shown in the heat map. Data in **Fig. S5 B and** Fig. 5 **D** are identical/from the same experiment. * p < 0.05, ** p < 0.01, *** p < 0.001, and **** p < 0.0001. (**C**) Balb/c mice were infected either by aerosol-inhalation with 3x LD50 PR8 or intranasal droplet challenge with 5x LD50 PR8. Lungs were removed four days later, and viral titers were measured via plaque assay. N=5 mice.; Mann-Whitney test was utilized for lung viral titers on the specified day. N.s. = not significant

**Table S1.** M2e-Consensus sequence (CS) alanine scanning peptide library.

| AA | 1 | 2 | 3 | 4 | 5 | 6 | 7 | 8 | 9 | 10 | 11 | 12 | 13 | 14 | 15 | 16 | 17 | 18 | 19 | 20 | 21 | 22 | 23 | Name |
| --- | --- | --- | --- | --- | --- | --- | --- | --- | --- | --- | --- | --- | --- | --- | --- | --- | --- | --- | --- | --- | --- | --- | --- | --- |
| M2e consensus | S | L | L | T | E | V | E | T | P | I | R | N | E | W | G | C | R | C | N | D | S | S | D | M2e-CS |
| Alanine Scanning Library | A | L | L | T | E | V | E | T | P | I | R | N | E | W | G | C | R | C | N | D | S | S | D | S1A |
|  | S | A | L | T | E | V | E | T | P | I | R | N | E | W | G | C | R | C | N | D | S | S | D | L2A |
|  | S | L | A | T | E | V | E | T | P | I | R | N | E | W | G | C | R | C | N | D | S | S | D | L3A |
|  | S | L | L | A | E | V | E | T | P | I | R | N | E | W | G | C | R | C | N | D | S | S | D | T4A |
|  | S | L | L | T | A | V | E | T | P | I | R | N | E | W | G | C | R | C | N | D | S | S | D | E5A |
|  | S | L | L | T | E | A | E | T | P | I | R | N | E | W | G | C | R | C | N | D | S | S | D | V6A |
|  | S | L | L | T | E | V | A | T | P | I | R | N | E | W | G | C | R | C | N | D | S | S | D | E7A |
|  | S | L | L | T | E | V | E | A | P | I | R | N | E | W | G | C | R | C | N | D | S | S | D | T8A |
|  | S | L | L | T | E | V | E | T | A | I | R | N | E | W | G | C | R | C | N | D | S | S | D | P9A |
|  | S | L | L | T | E | V | E | T | P | A | R | N | E | W | G | C | R | C | N | D | S | S | D | I10A |
|  | S | L | L | T | E | V | E | T | P | I | A | N | E | W | G | C | R | C | N | D | S | S | D | R11A |
|  | S | L | L | T | E | V | E | T | P | I | R | A | E | W | G | C | R | C | N | D | S | S | D | N12A |
|  | S | L | L | T | E | V | E | T | P | I | R | N | A | W | G | C | R | C | N | D | S | S | D | E13A |
|  | S | L | L | T | E | V | E | T | P | I | R | N | E | A | G | C | R | C | N | D | S | S | D | W14A |
|  | S | L | L | T | E | V | E | T | P | I | R | N | E | W | A | C | R | C | N | D | S | S | D | G15A |
|  | S | L | L | T | E | V | E | T | P | I | R | N | E | W | G | A | R | C | N | D | S | S | D | C16A |
|  | S | L | L | T | E | V | E | T | P | I | R | N | E | W | G | C | A | C | N | D | S | S | D | R17A |
|  | S | L | L | T | E | V | E | T | P | I | R | N | E | W | G | C | R | A | N | D | S | S | D | C18A |
|  | S | L | L | T | E | V | E | T | P | I | R | N | E | W | G | C | R | C | A | D | S | S | D | N19A |
|  | S | L | L | T | E | V | E | T | P | I | R | N | E | W | G | C | R | C | N | A | S | S | D | D20A |
|  | S | L | L | T | E | V | E | T | P | I | R | N | E | W | G | C | R | C | N | D | A | S | D | S21A |
|  | S | L | L | T | E | V | E | T | P | I | R | N | E | W | G | C | R | C | N | D | S | A | D | S22A |
|  | S | L | L | T | E | V | E | T | P | I | R | N | E | W | G | C | R | C | N | D | S | S | A | D23A |

**Table S2.**
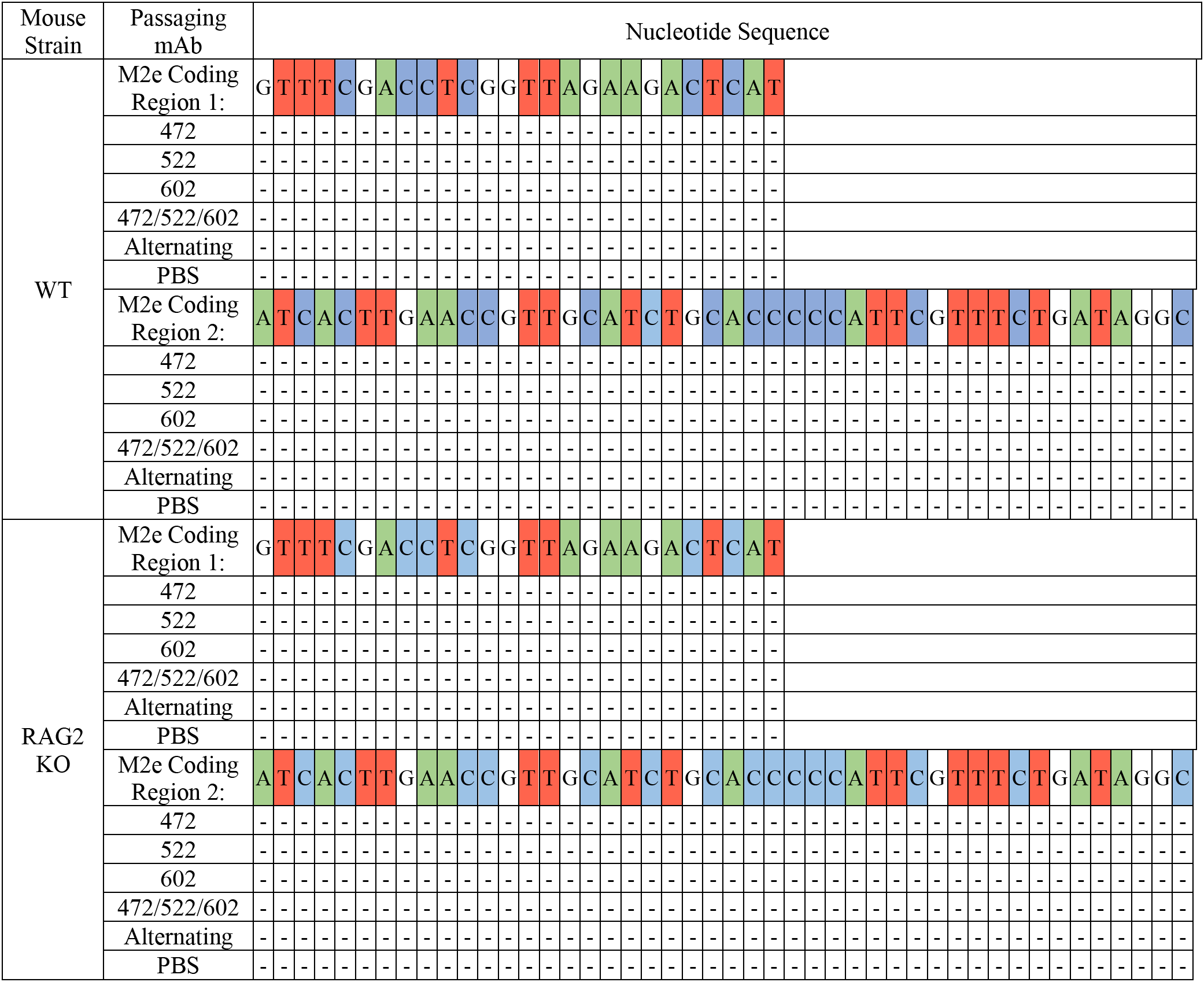
M2e sequences of influenza A viruses isolated from WT or RAG2-KO mice after single M2e-MAb or M2e-MAb cocktail antibody treatments. Chromatograms of M2e sequences for the M2e coding regions of viral isolates from passage experiments outlined in Fig. S2. Dash indicates sequence conservation

